# Resetting H3K4me3, H3K27ac, H3K9me3 and H3K27me3 during the maternal-to-zygotic transition and blastocyst lineage specification in bovine embryos

**DOI:** 10.1101/2022.04.07.486777

**Authors:** Chuan Zhou, Michelle M. Halstead, Amèlie Bonnet-Garnier, Richard M. Schultz, Pablo J. Ross

## Abstract

It remains poorly understood how histone modifications regulate changes in gene expression during preimplantation development. Using a bovine model, we profiled changes in two activating (H3K4me3 and H3K27ac) and two repressive (H3K9me3 and H3K27me3) marks in oocytes, 2-, 4- and 8-cell embryos (that developed in the presence or absence of the transcription inhibitor a-amanitin), morula, blastocysts, inner cell mass cells and trophectoderm. In oocytes, we find that broad bivalent domains of H3K4me3 and H3K27me3 mark developmental genes, and that prior to genome activation, H3K9me3 and H3K27me3 co-occupy gene bodies. During genome activation, chromatin accessibility is established before canonical H3K4me3 and H3K27ac, and although embryonic transcription is required for this active remodeling, it is dispensable for maintenance of pre-established histone marks. Finally, blastocyst lineages are defined by differential Polycomb repression and transcription factor activity. Overall, these results further support the use of bovine as a more appropriate model system than the mouse to study genome activation and cell lineage specification during human preimplantation development.

## Introduction

Preimplantation development encompasses two fundamentally important, but poorly understood, cell identity transitions in biology: the acquisition and subsequent loss of totipotency. Key to these transitions is the maternal-to-zygotic transition (MZT), marked by embryonic genome activation (EGA) and clearance of oocyte-specific messages ^1^. As blastomeres continue to divide, they progressively transition from totipotency to pluripotency, and finally segregate into two cell types that comprise the blastocyst: the inner cell mass (ICM) and trophectoderm (TE). Events such as the MZT and blastocyst formation correspond to both changes in gene expression and wide-spread epigenetic remodeling of DNA methylation, histone modifications and chromatin structure ^2^. Unclear, however, is the causal relationship between transcriptomic changes and chromatin remodeling.

Modification of histone tails, for example, can modulate gene expression by controlling the accessibility of chromatin to transcription factors (TFs) and RNA polymerase ^3–5^. Of many possible modifications of histone tail residues, a select few are commonly profiled to characterize active and repressed genomic regions. These include trimethylation of histone 3 lysine 4 (H3K4me3), which marks active promoters, acetylation of histone 3 lysine 27 (H3K27ac), which marks both active promoters and enhancers, trimethylation of histone 3 lysine 27 (H3K27me3), a mark of Polycomb repression and facultative heterochromatin, and trimethylation of histone 3 lysine 9 (H3K9me3), associated with constitutive heterochromatin.

Cross-talk between these modifications and their distribution throughout the genome collectively allows for fine-tuning of spatial and temporal gene expression. Unsurprisingly, disruption of this dynamic has catastrophic effects on transcription and development ^4^.

Although several recent reports have shed light on epigenetic remodeling during preimplantation development, they either do not capture the entire continuum of stages or focus on discrete levels of regulation, e.g., chromatin accessibility ^6–13^ or select histone modifications ^14–21^. Efforts to integrate such datasets post hoc can be problematic, because technical and methodological differences, such as missing biological timepoints and differing sample composition, can confound downstream analyses. Thus, the collective dynamics of histone modifications, chromatin structure, and DNA methylation during preimplantation development, and how these changes relate to changes in gene expression – and therefore cell identity and developmental potential – remain largely unknown, especially in mammals other than mouse. This gap in knowledge is especially conspicuous, because accumulating evidence suggests that larger mammals, e.g., cattle, constitute a better model system than rodents to understand human preimplantation development ^12,20^.

We report here a comprehensive analysis of both active (H3K4me3, H3K27ac) and repressive histone modifications (H3K9me3, H3K27me3) that bridges key developmental transitions from fertilization to blastocyst formation in bovine embryos. In addition, we integrate these results with previously published DNA methylation, chromatin accessibility, and gene expression datasets. We find a dramatic shift in the overall structure of the epigenome between the 8-cell and morula stages. Prior to EGA, embryos are distinguished by broad domains of all profiled histone marks, as well as enrichment for repressive marks in gene bodies. Post-EGA, this naïve chromatin organization resolves into a more typical structure, with chromatin accessibility appearing earlier than canonical H3K4me3 and H3K27ac. Inhibiting embryonic transcription severely impacts H3K27ac remodeling, leading to the retention of a 4-cell-like epigenetic state in 8-cell embryos. Finally, the two cell lineages in blastocysts — ICM and TE — are defined by accumulation of differential Polycomb repression and distinct regulatory circuitry, which was notably similar to that observed in human blastocysts, further supporting the use of bovine as a relevant model for human preimplantation development.

## Results

### Histone modification profiles during bovine preimplantation development

Using low-input CUT&Tag (cleavage under targets & tagmentation) ^22^, H3K4me3, H3K27ac, H3K9me3, and H3K27me3 were profiled in bovine germinal-vesicle (GV) oocytes, 2-, 4- and 8-cell embryos (that developed in the presence or absence of the transcription inhibitor a-amanitin), morula, blastocysts, ICM and TE (n=2 biological replicates). In total, 72 CUT&Tag libraries were generated and sequenced, producing 6,858,594,136 raw reads, from which 1,387,865,654 informative alignments were obtained and used for downstream analyses (**Extended Tables 1-4**). To complement data previously published by our group ^12^, chromatin accessibility was profiled in blastocysts, ICM, and TE (n=3 biological replicates) using ATAC-seq (assay for transposase-accessible chromatin by sequencing) ^23^ (**Extended Fig. 1a-b, Extended Table 5**). Genome-wide profiles of CUT&Tag biological replicates were highly correlated (average Pearson’s R 0.96) (**Extended Fig. 1c**). For each histone mark, samples clustered into two general groupings: pre-EGA stages (2-, 4-, and 8-cell embryos), and post-EGA stages (morula, blastocysts, ICM, and TE) (**Fig. 1a**). For each stage and mark, regions with significant enrichment in both biological replicates, e.g., peaks, were identified.

**Figure 1.**
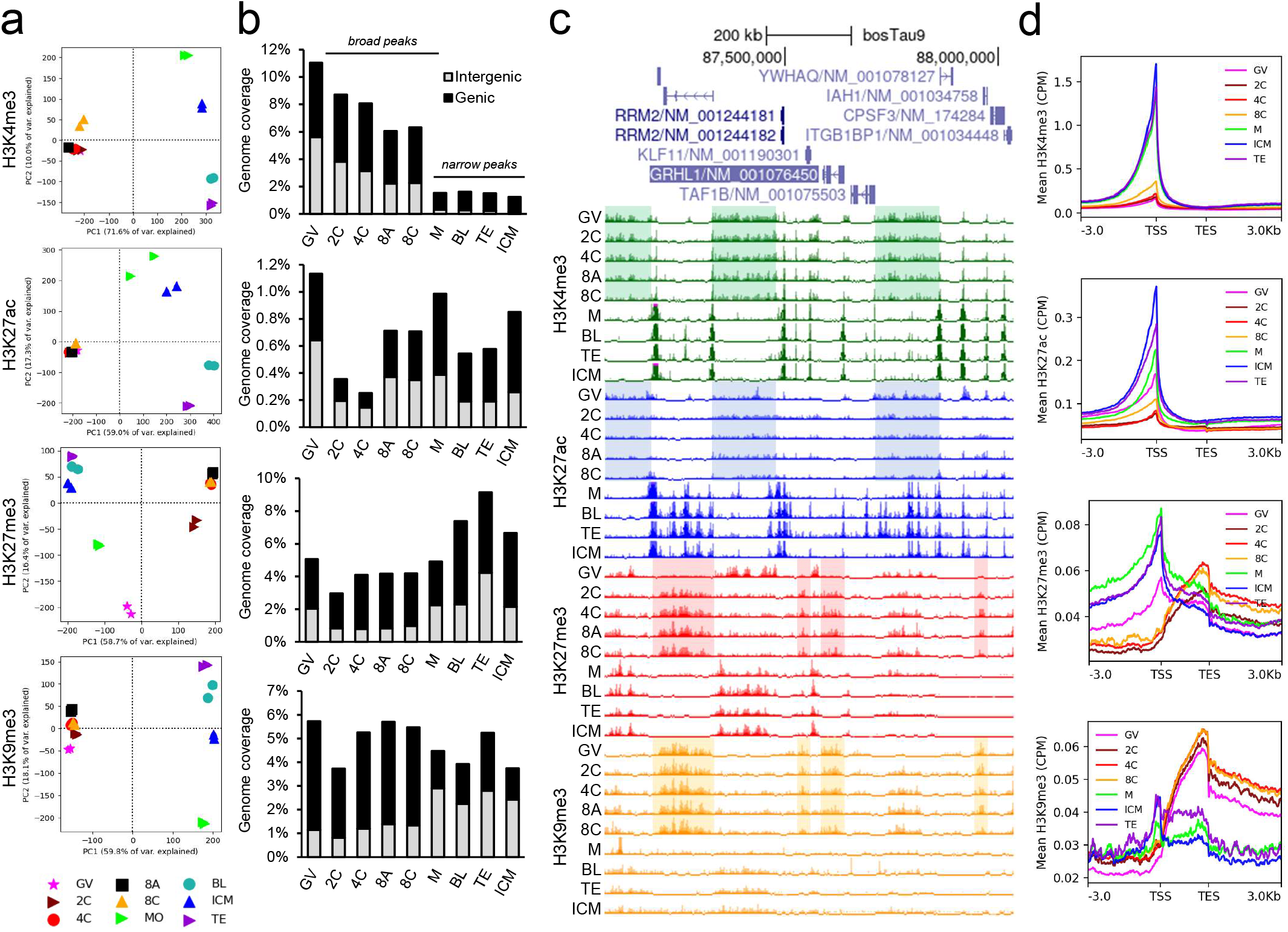
Profiles of histone modifications in bovine oocytes and preimplantation embryos. **a**, Principal components analysis (PCA) of CUT&Tag libraries. **b**, Genome coverage of peaks identified in both biological replicates. Peaks classified as genic (overlapping 2Kb promoter or gene bodies) or intergenic. For H3K4me3, genome coverage calculated for broad peaks until the 8-cell stage, then for narrow peaks. **c**, Normalized signal (counts per million; CPM) of one biological replicate per developmental stage and histone mark. Shaded regions correspond to noncanonical broad distributions (green, blue) and intragenic enrichment (red, orange). Viewing range from 0 to 1.5 CPM. Values exceeding maximum range indicated by pink bars. **d**, Average normalized signal (CPM) for each histone mark at transcription start sites (TSS) and end sites (TES). Transcripts scaled to 2Kb, 3Kb up- and downstream regions shown. GV oocytes (GV), 2-cell (2C), 4-cell (4C), 8-cell embryos (8C), a-amanitin-treated 8-cell embryos (8A), morula (M), blastocyst (BL), trophectoderm (TE) and inner cell mass (ICM).

Generally, H3K4me3 and H3K27ac were treated as “narrow” marks when calling peaks, and H3K27me3 and H3K9me3 as “broad” marks. However, considering recent findings that broad domains of noncanonical H3K4me3 (ncH3K4me3) are present in bovine oocytes ^20^, for peak calling purposes H3K4me3 was treated as a broad mark up until the 8-cell stage (see Methods).

Global abundance of a given histone modification, measured as percent of the genome covered by peaks, changed dramatically throughout the course of preimplantation development (**Fig. 1b**). In oocytes and pre-EGA embryos, H3K4me3 was wide-spread, covering 11% of the genome in GV oocytes (**Fig. 1b**). These broad domains of H3K4me3 were clearly evident up until the 8-cell stage, at which point they resolved into narrow peaks at promoters (**Fig. 1c**). H3K27ac also occupied broad domains in pre-EGA embryos, but the H3K27ac signal was generally weaker than that observed for H3K4me3 (**Fig. 1c**). Indeed, genomic coverage of H3K27ac decreased substantially after fertilization, with partial re-establishment in 8-cell embryos (**Fig. 1b**), although canonical narrow peaks of H3K27ac were not abundantly evident until the morula stage (**Fig. 1c**). Compared to these activating marks, H3K9me3 abundance was relatively stable throughout preimplantation development (**Fig. 1b**). However, the localization of this mark underwent a remarkable shift after fertilization, with H3K9me3 in oocytes and pre-EGA embryos occurring primarily at gene bodies (**Fig. 1c-d**), then switching to an intergenic distribution between the 8-cell and morula stages (**Fig. 1b**).

Similar to H3K9me3, H3K27me3 coverage was relatively constant in oocytes and pre-EGA embryos (**Fig. 1b**). However, unlike H3K9me3, the localization of this mark changed several times, from primarily intergenic in oocytes, to intragenic in pre-EGA embryos, and then back to intergenic after EGA (**Fig. 1c-d**). This profile contradicts immunofluorescence studies, which found that H3K27me3 is progressively and globally erased after fertilization, and almost absent in 8-cell embryos ^24–26^. In contrast to our findings, a recent study using CUT&RUN (cleavage under targets and release using nuclease) ^27^ indicated global loss of H3K27me3 following fertilization from the 4- to 16-cell stage ^20^. The likely explanation for this difference is that 10-fold fewer cells were used, noting that when similar numbers of cells were used, e.g., to profile blastocyst, a high degree of concordance was observed between CUT&Tag and CUT&RUN libraries (**Extended Fig. 1d**).

Overall, the maternal profiles of H3K4me3 and H3K9me3 were highly similar to that observed in 2-, 4-, and 8-cell embryos, suggesting substantial inheritance from the oocyte, both at intergenic regions marked by ncH3K4me3, and at gene bodies marked by H3K9me3. On the other hand, H3K27ac underwent substantial depletion after fertilization, and although the oocyte-specific H3K27me3 is largely erased in 2-cell embryos, *de novo* establishment of this mark was evident at gene bodies.

### Broad bivalent domains in oocytes mark developmental genes

The results described above raise the question about the relationship between remodeling of histone modifications and the repression and activation of gene expression programs that underlie changes in cell identity. Other studies have shown that the chromatin structure of pre-EGA embryos is markedly immature, characterized by high histone mobility ^28^, dispersed chromatin ^29^, and lack of canonical 3-D chromatin architecture ^30^. With developmental time, the chromatin adopts a more mature, condensed structure, at which point enhancer activity once again becomes necessary to relieve repression and activate of gene expression ^31^.

Consequently, the genomic distribution of histone modifications during preimplantation development, especially prior to EGA, does not necessarily follow canonical rules, and may not serve typical functions. For instance, H3K4me3, which generally occurs as sharp peaks at promoters in differentiated cells, occupies broad intergenic regions (> 5 Kb) in most mammalian oocytes and pre-EGA embryos ^14,20^. To account for this biology, H3K4me3 was treated as a “broad” mark for peak calling purposes, which improved reproducibility between biological replicates in GV oocytes, 2-, 4-, and 8-cell embryos (**Extended Fig. 2a**). Unexpectedly, treating H3K27ac as a broad mark also improved reproducibility at these stages (**Extended Fig. 2a**), indicating that this mark also has a broad noncanonical distribution prior to EGA.

Noncanonical H3K4me3 is anticorrelated with DNA methylation and occurs almost exclusively in partially methylated domains (PMDs) ^14,16,17,20^. To determine whether this pattern was recapitulated in bovine oocytes and embryos, and how PMDs relate to other histone modifications, we first identified PMDs in GV oocytes from published DNA methylation data ^32^ using methods similar to those described by Lu et al (2021) ^20^ (see Methods). We found that PMDs covered 25.3% of the genome, consistent with previous reports ^20^. Noncanonical H3K4me3 and H3K27ac were apparent within PMDs (**Extended Fig. 2b-c**). In GV oocytes, H3K4me3 was found almost exclusively in PMDs (90% of broad H3K4me3 peaks overlapped PMDs), although only 34% of PMDs were marked by H3K4me3 (**Extended Fig. 2d-e**). Enrichment of H3K4me3 in PMDs was maintained until the 8-cell stage. H3K27ac demonstrated a similar pattern. In GV oocytes, about 80% of broad H3K27ac domains occurred within PMDs, and this enrichment was maintained until the 4-cell stage (**Extended Fig. 2e**). H3K27me3 was also enriched at PMDs in GV oocytes, with more than 80% of H3K27me3 peaks overlapping PMDs. However, this enrichment was rapidly lost after fertilization, and H3K27me3 was not re-established at these regions until after EGA (**Extended Fig. 2e**). In contrast to the other three marks, H3K9me3 was notably depleted at PMDs in GV oocytes (only 10% of peaks overlapped with PMDs), and was gradually established in PMDs as development progressed (**Extended Fig. 2c-e**). This lack of H3K9me3 enrichment at PMDs was anticipated given global demethylation of the embryonic genome after fertilization ^32,33^, and that H3K9me deposition can depend on DNA methylation, and vice versa ^34^.

H3K4me3, H3K27me3, and H3K27ac were all enriched at PMDs. Because H3K27ac and H3K27me3 are mutually exclusive modifications, this enrichment suggested that different PMDs are probably marked by different combinations of histone modifications, which could entail distinct functions. To identify these patterns, PMDs were grouped according to histone modifications and chromatin accessibility signal in GV oocytes. The analysis revealed four clusters of PMDs (**Fig. 2a**), each exhibiting unique developmental changes (**Extended Fig. 3a**).

**Figure 2.**
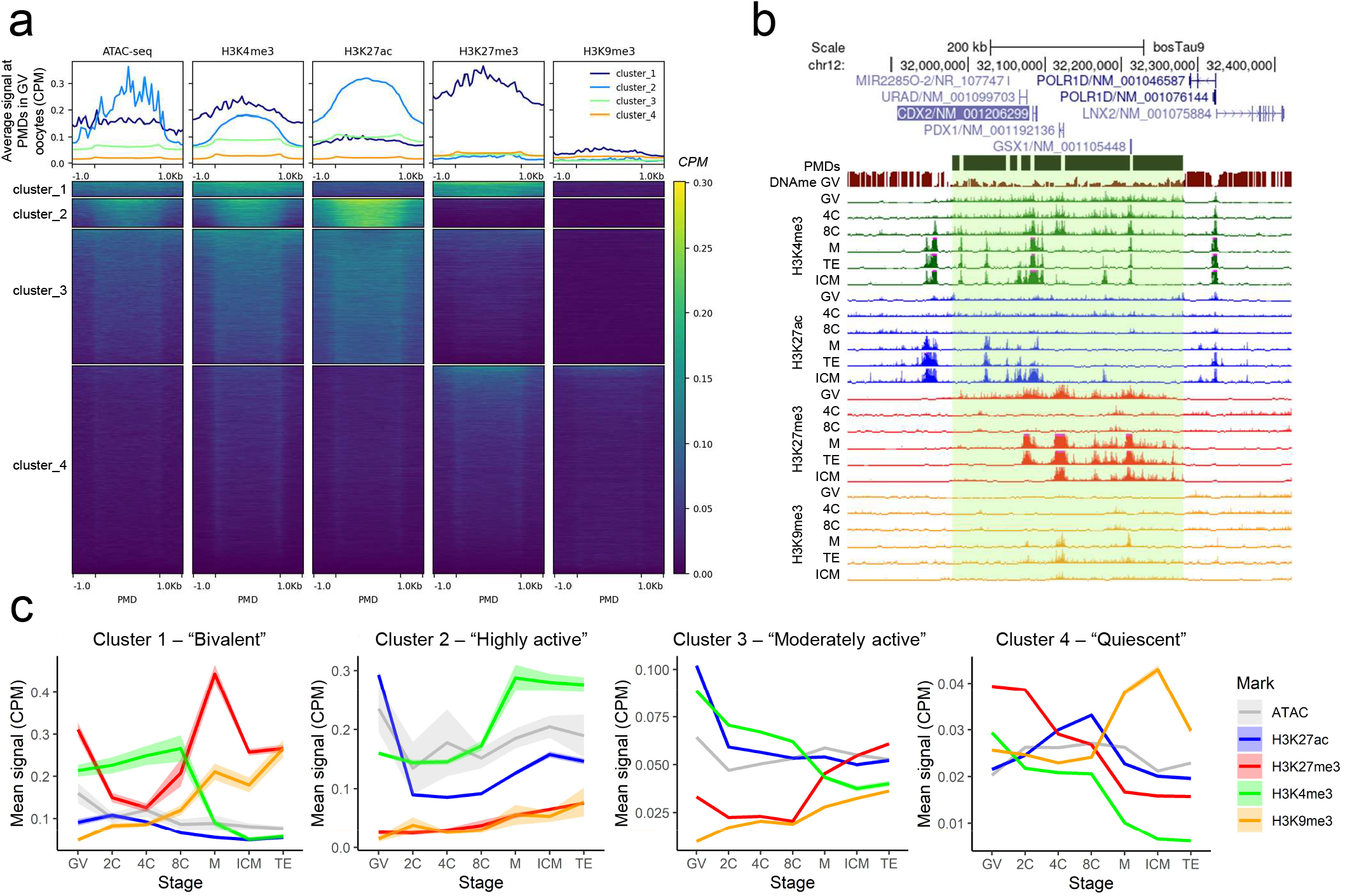
Characterization of partially methylated domains (PMDs) in GV oocytes. **a**, Clustering of PMDs based on normalized signal of H3K4me3, H3K27ac, H3K27me3, H3K9me3, and ATAC-seq in GV oocytes. Signal normalized by CPM in 100 bp windows, with PMDs scaled to 3Kb, and showing regions 1Kb up- and downstream. **b**, Representative gene track image of a bivalent PMD (“cluster_1”), overlapping three homeobox genes: *CDX2, GSX1*, and *PDX1*. One biological replicate shown per stage and mark. Viewing range from 0 to 1.5 CPM. Values exceeding maximum range indicated by pink bars. **c**, Average epigenetic signal (CPM ± standard error) for PMDs belonging to each cluster across development.

PMDs belonging to “cluster_1”, hereafter referred to as bivalent PMDs, were co-marked by H3K4me3 and H3K27me3 in GV oocytes (**Fig. 2a-b**), and covered 2% of the genome.

Although H3K4me3 was maintained at bivalent PMDs up to the 8-cell stage, H3K27me3 was rapidly erased following fertilization before re-establishment in morula (**Fig. 2c**). Co-occurrence of H3K4me3 and H3K27me3, termed “bivalency,” has been extensively described in embryonic stem cells (ESC), where bivalent domains tend to mark developmentally-related genes, poising them for future expression ^35^. There is little evidence for bivalency in rodent embryos prior to the blastocyst stage ^15,17^, but H3K4me3 and H3K27me3 are known to co-occur in PMDs in both porcine and bovine oocytes ^20^.

Specifically, we find that bivalent PMDs in bovine GV oocytes mark genes related to transcriptional activation and development, and especially conserved homeobox genes (**Extended Table 6**). Moreover, among all PMD clusters, genes marked by bivalent PMDs were the least expressed throughout preimplantation development (**Extended Fig. 3b**), suggesting a role in poising these genes for future expression, similar to the role of bivalency in ESC. Both “cluster_2” and “cluster_3” demonstrated enrichment for activating marks at early stages, but “cluster_2” PMDs, hereafter referred to as highly active PMDs, were strongly and briefly enriched for H3K27ac, demonstrated higher accessibility in GV oocytes, and marked genes related to transcription silencing (**Fig. 2c, Extended Table 6**). These strongly active PMDs might therefore play a role in maintaining transcriptional quiescence in full-grown oocytes ^1^. On the other hand, moderately active PMDs (“cluster_3”), were not as enriched for activating marks as strongly active PMDs, and rather than occurring near genes related to transcriptional repression, instead occurred near genes related to transcription activation (**Fig. 2c, Extended Table 6**). Finally, the most abundant category of PMDs (“cluster_4,” termed quiescent PMDs), demonstrated low enrichment for most marks, with slightly higher levels of H3K27me3 in GV oocytes (**Fig. 2a,c**). Genes marked by these regions demonstrated little functional enrichment (**Extended Table 6**). As such, these PMDs likely represent heterochromatin compartments in differentiated cells, especially given the accumulation of H3K9me3 during later stages of development (**Fig. 2c**), which was also maintained in fetal fibroblasts (**Extended Fig. 3c**).

The functional significance of PMDs for bovine preimplantation embryos is unknown. In mouse, improper retention of ncH3K4me3 through knock down of lysine demethylases (KDM5A and KDM5B) leads to downregulation of EGA-related genes and impaired blastocyst formation ^14,15^, suggesting a role preparing the genome for transcriptional activation. Conversely, overexpression of KDM5B leads to the removal of ncH3K4me3 and improper gene silencing in oocytes. Thus, ncH3K4me3 may also play a silencing role, at least during oogenesis, perhaps by acting like a “molecular sponge” to reduce the effective concentration of TFs, and thereby their ability to induce transcription ^16^. In particular, the functional significance of bivalent domains in bovine oocytes remains to be determined, noting that there is little evidence of bivalency at PMDs in mouse oocytes ^14,16,17,20,36^. In summary, these broad domains, devoid of DNA methylation and precisely marked by specific combinations of histone modifications, likely comprise a unique mechanism to both repress transcription and poise developmental genes for future expression.

### H3K9me3 and H3K27me3 co-occupy gene bodies in pre-EGA embryos

Given the extensive changes to chromatin structure at intergenic regions, we next focused on the genes themselves. H3K4me3 and H3K27ac signal at promoters increased progressively after fertilization, whereas H3K27me3 and H3K9me3 underwent a remarkable shift in distribution from gene bodies (“intragenic”) in 2-, 4-, and 8-cell embryos, to promoters from the morula stage onward (**Fig. 1d**). Notably, intragenic H3K9me3 was already evident in GV oocytes, but intragenic H3K27me3 was not, indicating that maternal H3K9me3 is inherited, whereas intragenic H3K27me3 is deposited *de novo* following fertilization. To determine if this pattern was characteristic of all genes or only a subset, transcripts were clustered based on their H3K27me3 and H3K9me3 signatures (**Extended Fig. 4a**).

A small subset of transcripts (“cluster_1”; n=1,639) were depleted for both repressive marks in pre-EGA embryos, but strongly marked from the morula stage onward; these transcripts also exhibited strong H3K27me3 signal in GV oocytes, which was promptly erased after fertilization (**Fig. 3a-b, Extended Fig. 5a**). Moreover, they were broadly marked by H3K4me3 and H3K27ac, rather than demonstrating sharp peaks at their transcription start sites (TSS), (**Fig. 3a, Extended Fig. 4b**), and were expressed at lower levels than genes belonging to the other two clusters (**Extended Fig. 4c**). As was the case for genes marked by bivalent PMDs, which demonstrated similar patterns of histone remodeling, homeobox genes were highly overrepresented in “cluster_1” (**Extended Fig. 5b**). Therefore, “cluster_1” genes are hereafter referred to as bivalent genes. Notably, not all homeobox genes belonged to this cluster. For instance, several homeobox genes implicated in early development and pluripotency (e.g., *NANOG, POU5F1, DUX4*), corresponded instead to “cluster_3.”

**Figure 3.**
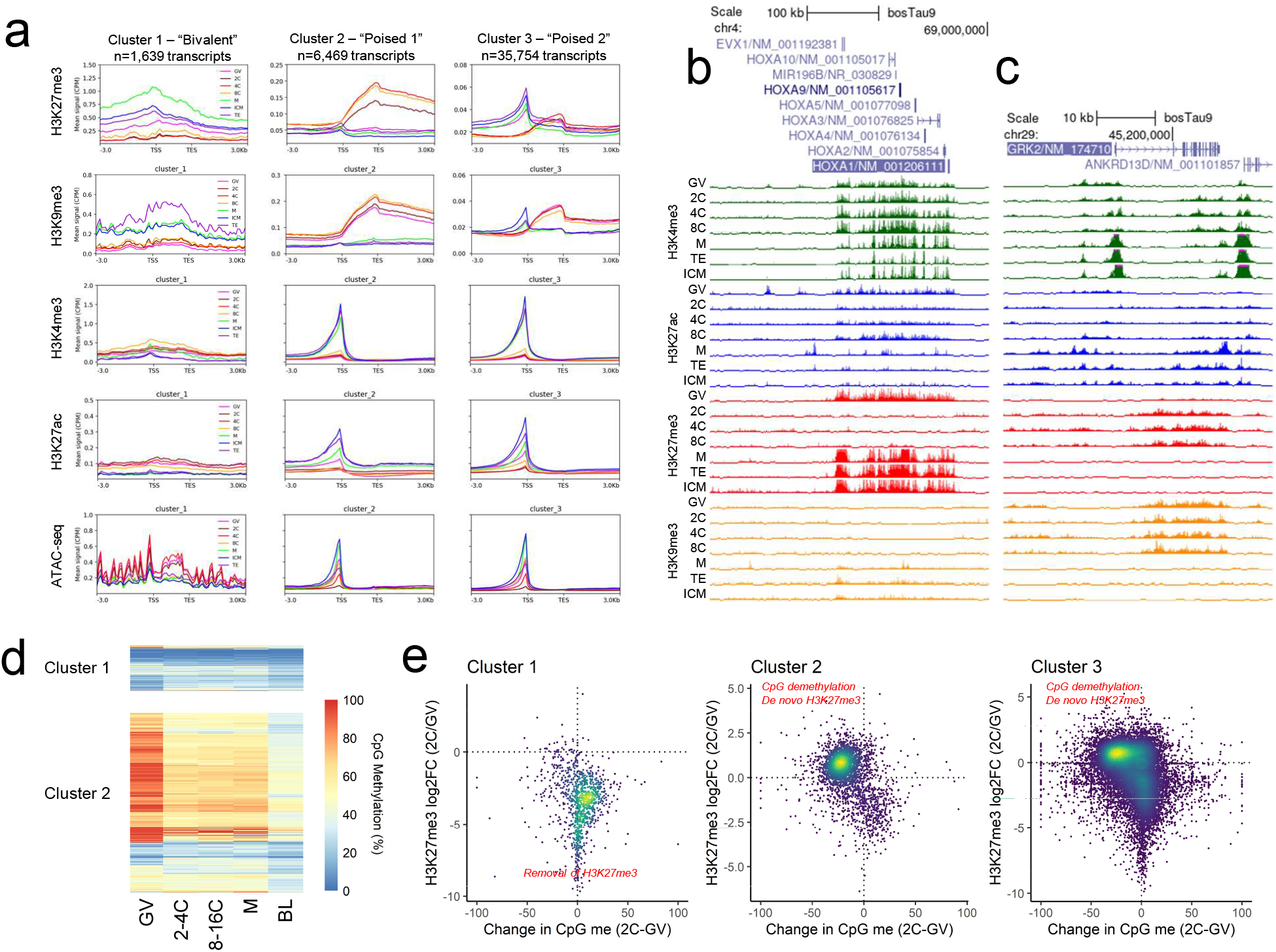
H3K9me3 and H3K27me3 mark gene bodies in pre-EGA embryos. **a**, Average normalized signal (CPM) of histone marks and chromatin accessibility for each transcript cluster, defined by kmeans clustering (k=3) based on normalized H3K27me3 and H3K9me3 signal (CPM) across development. Transcripts scaled to 2Kb, regions 3Kb up- and downstream shown. **b**, An example of “cluster_1” transcripts, the *HOX* cluster. **c**, An example of “cluster_2” transcripts: *GRK2* (a kinase). One biological replicate shown per stage and mark. Viewing range from 0 to 1.5 CPM. Values exceeding maximum range indicated by pink bars. **d**, Gene body CpG methylation (%) of “cluster_1” and “cluster_2” genes throughout development. **e**, For each group of transcripts, comparison of change in gene body CpG methylation (GV oocytes versus 2-to-4-cell embryos) to change in intragenic H3K27me3 signal (GV oocytes versus 2-cell embryos). Overall trends are summarized in red text.

The majority of transcripts belonged to either “cluster_2” (n=6,469) or “cluster_3” (n=35,754), both of which demonstrated intragenic H3K9me3 and H3K27me3 in pre-EGA embryos, although only intragenic H3K9me3 was inherited from the oocyte (**Fig 3a,c, Extended Fig. 4a**). In contrast to bivalent genes, the TSS of these transcripts were markedly enriched for H3K4me3, H3K27ac, and accessibility, albeit with varying intensity depending on the developmental stage (**Fig. 3a, Extended Fig. 4b**). Consequently “cluster_2” and “cluster_3” transcripts are hereafter referred to as “poised-1” and “poised-2” transcripts, respectively. “Poised-1” transcripts exhibited more intense intragenic H3K9me3 and H3K27me3, and were functionally enriched for ATP-binding and kinase activity (**Extended Fig. 5c-d**), whereas “poised-2” transcripts demonstrated subtler intragenic H3K9me3 and H3K27me3 and were enriched for functions related to transcript processing, such as poly(A) binding, ribosomes, and spliceosomes (**Extended Fig. 5e-f**).

The co-occurrence of these two marks in gene bodies is a phenomenon not previously described in embryos, and its functional significance in the context of development is therefore unknown. H3K9me3 and H3K27me3 are usually mutually exclusive, but they can co-occur at developmentally repressed genes in a variety of cell types, including stem cells ^37^ and extra-embryonic lineages ^38^. We postulated that H3K9me3 and H3K27me3 might serve a similar function in pre-EGA embryos by preventing the expression of oocyte-specific programs. However, there was no notable over-representation of maternal, minor EGA, or major EGA genes in any of the three clusters (**Extended Fig. 6**).

Consequently, we hypothesize that the presence of repressive histone marks in gene bodies is likely not a mechanism to repress maternal programs, nor to activate embryonic ones, but rather a means to globally suppress transcription of the embryonic genome prior to EGA. In GV oocytes, most genes (poised transcripts) were marked by high gene body H3K9me3, but still retained some H3K4me3 at their promoters (**Fig. 3a**). This situation is highly reminiscent of the poised state of lineage-specific genes in mesenchymal stem cells and preadipocytes, in which promoters are marked by H3K4me3 and gene bodies are marked by DNA methylation, which recruits SETDB1, leading to the deposition of H3K9me3 downstream of TSS ^39^. Intragenic H3K9me3 then prevents histone acetylation and spreading of H3K4me3, resulting in pausing of RNA polymerase and very low expression ^39^. Consistent with the poised state observed in these cells, genes with the highest levels of intragenic H3K9me3 in GV oocytes (“poised-1” transcripts) demonstrated significantly higher gene body methylation than those lacking intragenic H3K9me3 in GV oocytes (“bivalent” transcripts) (unpaired Wilcoxon signed-rank test; *p*<2.2e-16) (**Fig. 3d, Extended Fig. 4d**). Thus, in conjunction with DNA methylation, intragenic H3K9me3 may help to prevent aberrant gene transcription in the transcriptionally quiescent oocyte. After fertilization, however, the embryonic genome undergoes global DNA demethylation ^32,33^, including at gene bodies. For both “poised-1” and “poised-2” transcripts, methylation levels dropped after fertilization, and declined further upon blastocyst formation (**Extended Fig. 4d**). This loss of gene body methylation was concomitant with *de novo* deposition of intragenic H3K27me3 (**Fig. 3e**). The exchange of gene body methylation for H3K27me3 is not surprising, given that DNA methylation antagonizes deposition of H3K27me3 ^40^.

Overall, the presence of intragenic H3K9me3 in oocytes suggests a mechanism for general transcriptional repression until major EGA. As *de novo* deposition of intragenic H3K27me3 coincides with the loss of gene body methylation, this mark may serve a compensatory repressive role during global DNA demethylation, thus preventing premature transcription of the embryonic genome before the epigenome has been sufficiently reprogrammed to permit appropriate expression of developmental programs. Such a mechanism is reminiscent of H3K27me3 mediated noncanonical imprinting in mice, wherein H3K27me3 is temporarily inherited from the oocyte, leading to allele-specific repression of certain genes and paternal-specific X inactivation ^41^.

### Chromatin accessibility precedes establishment of canonical H3K4me3 and H3K27ac

Whereas all profiled histone marks underwent the greatest change in distribution and abundance during the 8-cell to morula transition (**Fig 1a-c**), the most dramatic change in chromatin accessibility occurred between the 4- and 8-cell stage (**Extended Fig. 1a**), when more than 68,000 ATAC-seq peaks were gained. This timing difference could reflect a priming mechanism, wherein a region becomes accessible before accumulating activating histone marks. Indeed, H3K27ac peaks that were established at the morula stage were already accessible in 8-cell embryos (**Fig. 4a**). The “accessibility first” pattern was even more dramatic for transcription start sites (TSS), which often gained accessibility as early as the 4-cell stage, but did not demonstrate canonical narrow H3K4me3 peaks until the morula stage (**Fig. 4b-c**).

**Figure 4.**
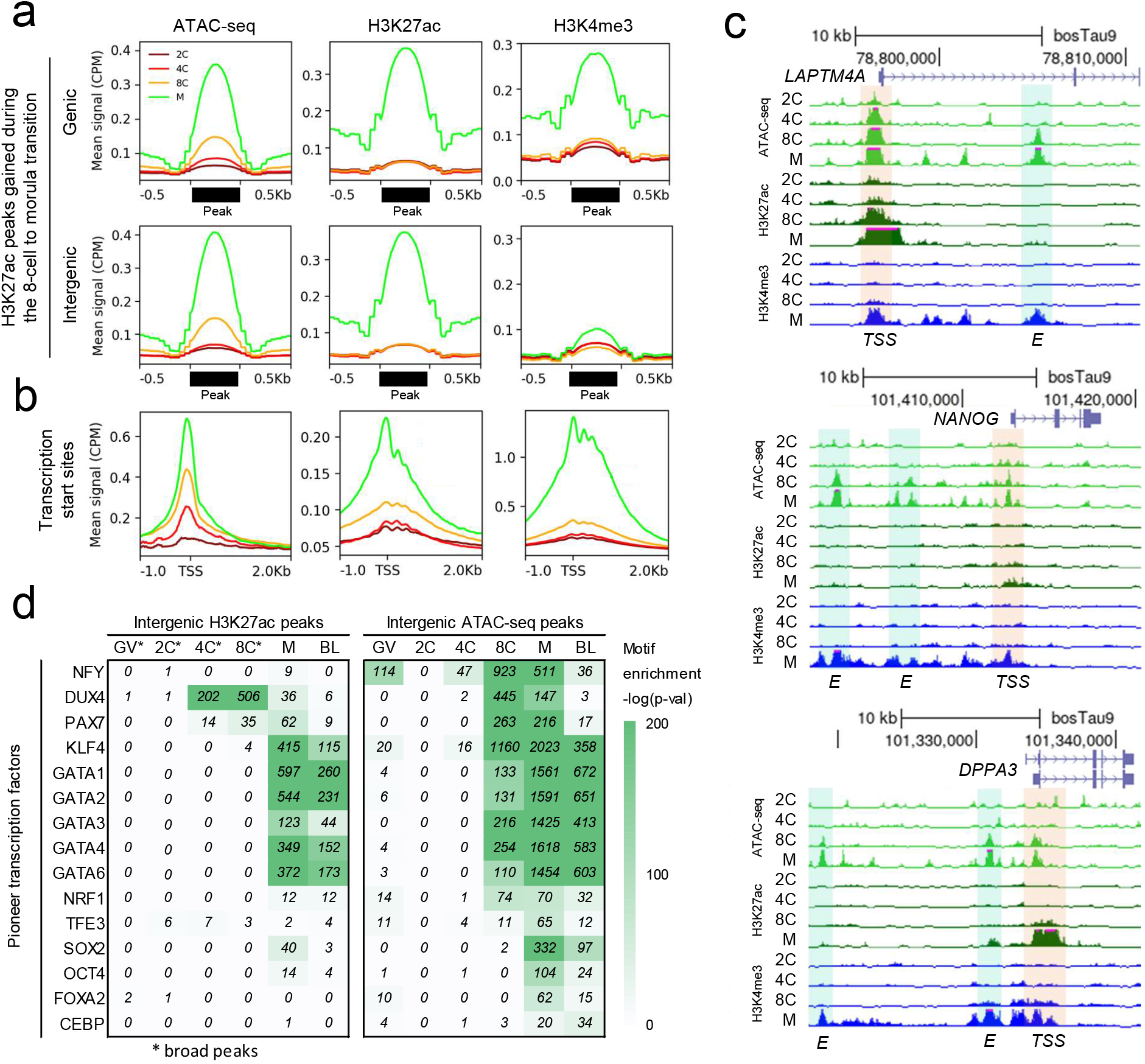
Chromatin accessibility precedes establishment of canonical H3K4me3 and H3K27ac in morula. **a**, Average ATAC-seq, H3K27ac, and H3K4me3 signal (CPM) at H3K27ac peaks which appeared during the 8-cell to morula transition. Peaks classified as genic if they overlapped gene bodies or promoters (2Kb upstream of TSS). **b**, Average signal at TSS. **c**, Normalized ATAC-seq, H3K27ac, and H3K4me3 signal for one biological replicate per developmental stage at the *LAPTM4A, NANOG*, and *DPPA3* loci. Shaded regions correspond to putative enhancers (E; blue) and promoters (TSS; orange). Viewing range from 0 to 1.5 CPM. Values exceeding maximum range indicated by pink bars. **c**, Enrichment of pioneer transcription factor motifs in H3K27ac and ATAC-seq peaks at each stage. From the GV to 8-cell stage, broad H3K27ac peaks (*) were used for motif enrichment analysis, then narrow peaks were used.

It seems unlikely that chromatin accessibility, in the absence of other mechanisms, would be the first epigenetic mark to establish a canonical structure after fertilization, as it is intimately linked to chromatin remodeling. For example, monomethylation of histone 3 lysine 4 (H3K4me1) can recruit chromatin remodeling complexes (e.g., SWI/SNF complex) leading to changes in accessibility ^42^. In fact, at developmental enhancers in mouse and human ESC, H3K4me1 often precedes nucleosomal depletion, H3K27ac, and enhancer activation ^43–45^. Moreover, in mouse, H3K4me1 is required for minor EGA ^46^. Thus, an unidentified step, possibly another histone modification, may precede establishment of accessible sites in 4- and 8-cell embryos, with later modifications (e.g., H3K27ac) serving to reinforce that accessibility.

Alternatively, but not mutually exclusive, is the possibility that pioneer TFs (TFs) are responsible, given their ability to bind to inaccessible chromatin and subsequently recruit remodeling complexes, histone modifiers, and other TFs ^47^. To address this hypothesis, motif enrichment analyses were conducted on intergenic ATAC-seq and H3K27ac peaks to determine whether the motifs of certain pioneer factors were over-represented in active regions prior to EGA. Although there was little enrichment for pioneer factor motifs in open chromatin at the 4-cell stage, H3K27ac peaks were already significantly enriched for DUX4 motifs (**Fig. 4d**) furthering the hypothesis that this TF can bind initially inaccessible regions and induce H3K27ac by recruiting histone acetyltransferases ^48^. By the 8-cell stage, DUX4 motifs were also enriched in open chromatin, along with the motifs of several other pioneer factors, including NFY, PAX7, KLF4, and GATA factors (**Fig. 4d**). Of note, whereas DUX4 motifs first appeared in H3K27ac peaks and then in accessible regions, the opposite was true for NFY, PAX7, KLF4, and GATA factor motifs, which first appeared in open chromatin, and then in H3K27ac peaks. The latter pattern – “accessibility first” – is consistent with the pre-establishment of accessibility at sites of future H3K27ac peaks (**Fig. 4a**), suggesting that pioneer factors contribute to the establishment of chromatin accessibility at the 8-cell stage prior to deposition of H3K27ac at these sites in morula.

Clearly, resolution of H3K4me3 and H3K27ac from broad domains to narrow peaks occurs only after chromatin condensation has been initiated, allowing for detection of discrete regions of accessibility. The identity of the factors that drive this transition (e.g., histone modifiers or pioneer TFs) remains unknown, and are the subject of ongoing investigation.

### Transcription is required for histone remodeling but not maintenance

As described above, temporal changes to histone modification distribution during preimplantation development occur on both a large scale (e.g., at PMDs) and at specific loci (e.g., genes and enhancers). The identity of factors responsible for these changes, and whether they are maternally-derived and/or products of embryonic transcription, remains unknown. To assess the contribution of embryonic transcription to epigenetic reprogramming, histone modifications were profiled in 8-cell embryos cultured from the 1-cell stage in the presence of a-amanitin. We focused on regions that gained, lost, or maintained enrichment (e.g., peaks) for a given histone modification during the 4- to 8-cell transition, and whether these regions corresponded to peaks in transcription-inhibited 8-cell embryos. Among H3K4me3, H3K27ac, H3K27me3, and H3K9me3, 73, 77, 52, and 63% of gained peaks were not established, respectively, and 36, 73, 50, and 59% of 4-cell peaks normally erased by the 8-cell stage were still present in 8-cell embryos treated with a-amanitin, respectively. In contrast, for all marks, about 90% of peaks found in both 4- and 8-cell embryos were also present in transcription-inhibited 8-cell embryos (**Extended Table 7**). Signal at these gained, lost, and retained peaks revealed that the epigenetic landscape of transcription-inhibited 8-cell embryos more closely resembled that of 4-cell embryos than 8-cell embryos (**Fig. 5a**), particularly for H3K27ac, which gained and lost the most peaks of any mark during the 4- to 8-cell transition (**Extended Table 7**).

**Figure 5.**
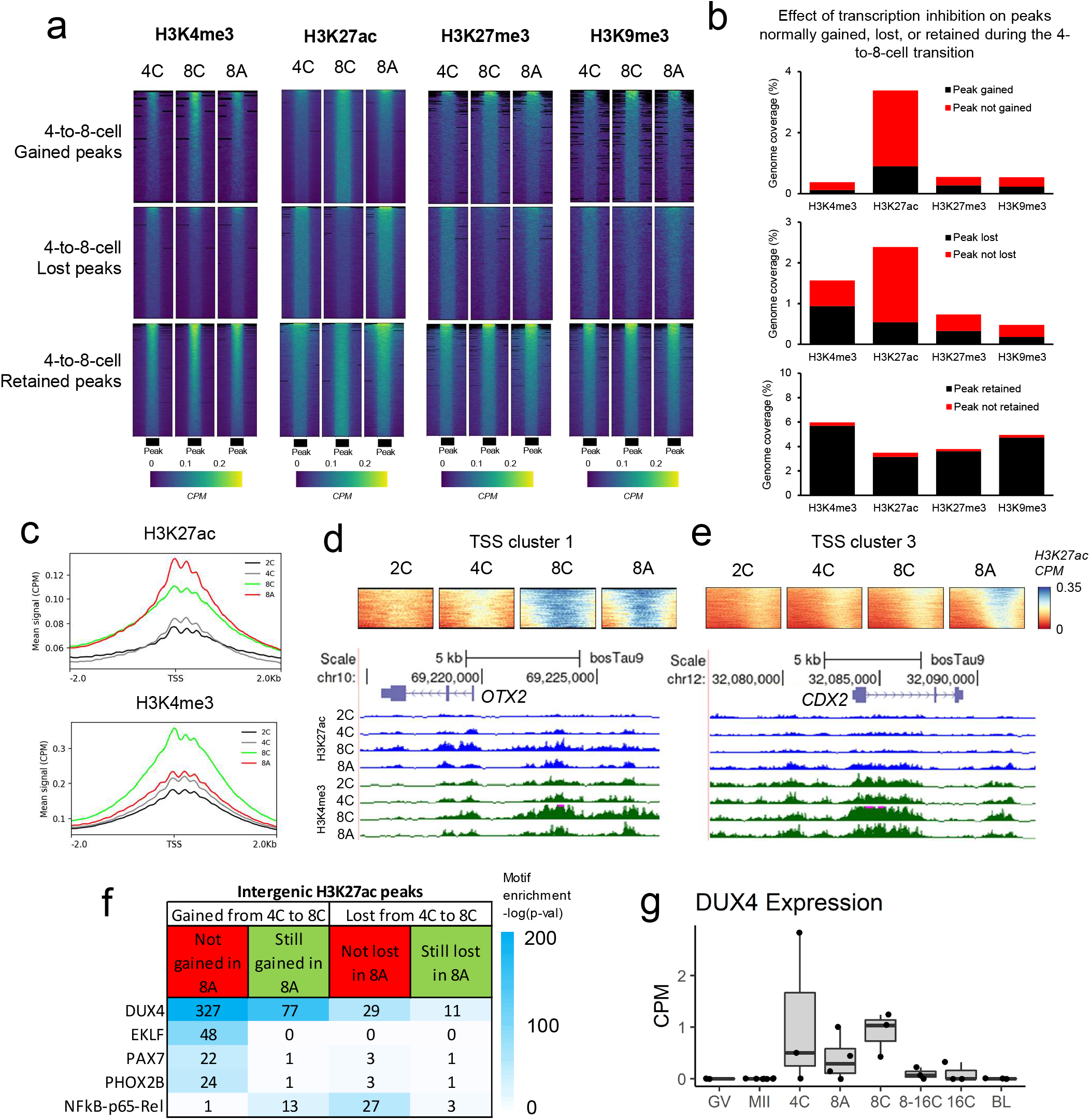
Effect of transcription inhibition on the epigenetic profile of 8-cell embryos. a, Normalized signal (CPM) in 4-cell (4C), 8-cell (8C) and 8-cell embryos cultured in the presence of a-amanitin (8A) at regions that gained, lost, or retained peaks during the 4- to 8-cell transition. **b**, Genomic coverage of regions that gained, lost, or retained peaks. Shading reflects the impact of transcription inhibition on peak status. Red indicates regions with a change in peak status in 8A embryos compared to 8C. **c**, Average signal (CPM) of active epigenetic marks at TSS in 2-cell (2C), 4C, 8C and 8A embryos at TSS. **d**, Normalized H3K27ac signal at “cluster 1” TSS, identified based on H3K27ac signal at TSS in 2C, 4C, 8C, and 8A embryos (Extended Fig. 17), and a gene track view of representative locus, *OTX2*. **e**, Normalized H3K27ac signal (CPM) at “cluster 3” TSS, and a gene track view of a representative locus, *CDX2*. **f**, Enrichment of select transcription factor motifs in intergenic H3K27ac peaks that were gained or lost during the 4- to 8-cell transition, and which were either sensitive or robust to inhibition of embryonic transcription. **g**, Normalized expression of *DUX4* (ENSBTAG00000049205) across development. Boxplots indicate the median and interquartile range (IQR), and whiskers span 1.5 times the IQR.

This disruption of histone remodeling is mirrored by the a-amanitin-sensitive expression of the respective histone modifiers. For instance, erasers of H3K4me3 were highly expressed in 8-cell embryos relative to oocytes and transcription-inhibited embryos (**Extended Fig. 7a**). In particular, *KDM5B*, which removes ncH3K4me3 in mouse embryos ^14–16^, was 30-fold more expressed in control relative to transcriptionally-inhibited 8-cell embryos. The time it would take to synthesize sufficient amounts of KDM5B protein from zygotically generated *KDM5B* transcripts may account for the observation that ncH3K4me3 is still present at the 8-cell stage (**Extended Fig. 5**). Regarding H3K27me3 modifiers, components of both Polycomb repressive complexes (PRC1 and 2) and the H3K27me3 demethylase *KDM6A* are zygotically-expressed in 8-cell embryos (**Extended Fig. 7b**). This expression could explain the erasure of intragenic H3K27me3 and establishment of intergenic H3K27me3 domains during the 8-cell to morula transition. In contrast, transcript abundance of H3K9me3 remodelers was generally insensitive to transcription inhibition, although *CBX3/HP1-gamma*, which binds to H3K9me3 and excludes H3K27me3 from the same loci ^49^, was upregulated in 8-cell embryos (**Extended Fig. 7c**), again reflecting the imminent loss of co-occupancy of H3K9me3 and H3K27me3 after the 8-cell stage. Zygotic expression of these key histone modifiers likely underpins the transition from a pre-EGA state, characterized by ncH3K4me3 and intragenic H3K9me3 and H3K27me3, to a post-EGA state, characterized by canonical distributions of these marks.

In contrast to H3K4me3, H3K9me3, and H3K27me3, the distribution of H3K27ac, similar to chromatin accessibility ^12^, was markedly perturbed in transcriptionally-inhibited 8-cell embryos. Transcription inhibition prevented 74% of regions that gained H3K27ac during the 4- to 8-cell transition from becoming marked, and resulted in incorrect retention of H3K27ac at 78% of regions that should have been deacetylated (**Fig. 5b**). These failures to deposit and remove H3K27ac at pertinent loci were reflected by downregulation of both histone acetyltransferases (*p300, PCAF*, and *KAT14*) and deacetylases (*HDAC1* and *HDAC2*) in response to a-amanitin treatment (**Extended Fig. 7d**). Moreover, 6,223 regions accumulated aberrant H3K27ac signal not present in either 4- or 8-cell embryos (**Extended Fig. 7e**), suggesting that factors directing HATs to target loci are also dysregulated. This aberrant H3K27ac signal was especially evident at TSS, despite reduced H3K4me3 (**Fig. 5c**).

Clustering of TSS based on H3K27ac signal revealed that transcription inhibition had the greatest impact on acetylation of a subgroup of transcripts (cluster 3; n=4,137 transcripts), with H3K27ac signal substantially increased downstream of their TSS (**Extended Fig. 7f**). This group was again enriched for homeobox genes (**Extended Table 8**), e.g., *CDX2* (**Fig. 5e**). Of note, another transcript cluster with high H3K27ac signal in both control and transcriptionally-inhibited 8-cell embryos (cluster 1; n=1,324 transcripts) was also enriched for homeobox genes (**Extended Table 8**), e.g., *OTX2* (**Fig. 5d**). These two distinct patterns at homeobox genes suggest separate modes of regulation for different subsets of developmentally-related genes. Overall, a significant portion of transcripts (clusters 1, 3, and 5) demonstrated increased H3K27ac signal at their TSS in 8-cell embryos that developed in the presence of a-amanitin. These genes were functionally enriched for processes involved in transcription regulation (**Extended Table 8**). This generalized increase in H3K27ac at promoters, which is normally a hallmark of transcriptional activation, is puzzling following a-amanitin treatment. However, because a-amanitin treatment prevents transcription elongation, but not binding of RNA polymerase II, polymerase stalling at TSS may be a contributing factor.

Regarding regions that were normally acetylated during the 4- to 8-cell transition, those which gained H3K27ac peaks regardless of a-amanitin treatment were primarily genic (68%) (**Extended Fig. 7g**), and marked genes related to cell division, e.g., actin and microtubule binding (**Extended Table 9**). In contrast, regions that failed to gain H3K27ac in transcriptionally inhibited embryos marked genes related to RNA and chromatin binding (**Extended Table 9**), and were more often intergenic (57%) (**Extended Fig. 7g**). H3K27ac peaks lost during this transition, regardless of whether they were also erased in transcription inhibited embryos, primarily marked genes involved in calcium signaling (**Extended Table 9**). Overall, these results suggest that H3K27ac remodeling mediated by maternal factors acts to (1) repress transcriptional programs related to oogenesis and fertilization, and (2) reinforce transcriptional programs related to cleavage. On the other hand, H3K27ac remodeling that depends on embryonic transcription seems to regulate programs related to epigenetic regulation and EGA.

Differential motif enrichment was also evident in these different sets of H3K27ac peaks. Peaks gained from the 4- to 8-cell stage that were dependent on embryonic transcription were enriched for KLF, PAX7, and PHOX2B motifs, which were not found in peaks gained regardless of a-amanitin treatment, nor in peaks lost during this transition (**Fig. 5f**). The opposite pattern was true for NFkB factors, which can participate in cross-talk with chromatin remodelers, e.g., HAT p300, and acetylation readers, e.g., Brd4 ^50,51^. NFkB motifs were uniquely present in gained H3K27ac peaks that did not depend on embryonic transcription, as well as regions that incorrectly retained H3K27ac when transcription was inhibited (**Fig. 5f**). As NFkB factors are maternally provided in mice ^52^ and transcripts are present in bovine oocytes ^12^, these results suggest dysregulation of a maternal system that directs HATs to target loci. Notably, in all peak sets, the top enriched motif was that of DUX4 (**Fig. 5f**), which is a pioneer factor that induces acetylation by recruiting the HAT p300, and which is increasingly implicated in the establishment of totipotency ^53,54^. The DUX4 motif was even enriched in peaks that were gained regardless of a-amanitin treatment (**Fig. 5f**), suggesting that functional DUX4 is present in transcriptionally inhibited embryos. Indeed, DUX4 appears to escape transcription inhibition (**Fig. 5g**). Because DUX4 is one of only a few genes expressed during minor EGA (**Extended Fig. 6c**), and RNA-seq data are unavailable for embryos before the 4-cell stage, DUX4 expression may have initiated before embryos were transferred to culture medium containing a-amanitin. Zygotic expression of DUX4 would be consistent with preliminary studies of human embryos, which indicate that DUX4 expression initiates during the 1-cell stage ^55^.

In summary, the impact of transcription inhibition on the epigenome is most severe for marks that began to resolve earliest, namely, H3K27ac. For marks that do not resolve to a canonical form until after the 8-cell stage (e.g., H3K4me3, H3K27me3, and H3K9me3), the impact of transcription inhibition was less evident at the 8-cell stage. The lack of embryonic expression of the appropriate modifiers, and therefore the inability to resolve the epigenome to a post-EGA state, may underlie the failure of a-amanitin treated embryos to progress past the 8-to-16-cell stage ^56^.

### Blastocyst lineages are defined by differential Polycomb repression

Following EGA, blastomeres progressively transition from totipotency to pluripotency, and by the blastocyst stage two distinct cell lineages are present: the ICM, a pluripotent lineage that forms the embryo proper, and the TE, a differentiated cell type that contributes to extra-embryonic structures (for detailed reviews, see ^57,58^). TE specification initiates in morula via a conserved mechanism in which outer blastomeres express GATA3, which upregulates TE-specific programs ^59^ (**Fig. 6a**). TE specification progresses more quickly in mouse, as the TE-specific factor CDX2 is already expressed in the outer cells of the morula, whereas CDX2 is not detected until blastocyst formation in human and cattle (**Fig. 6b**). Moreover, OCT4/POU5F1 expression remains widespread in human and bovine blastocysts, but is already restricted to the ICM in mouse ^58,60^ (**Fig. 6b**).

**Figure 6.**
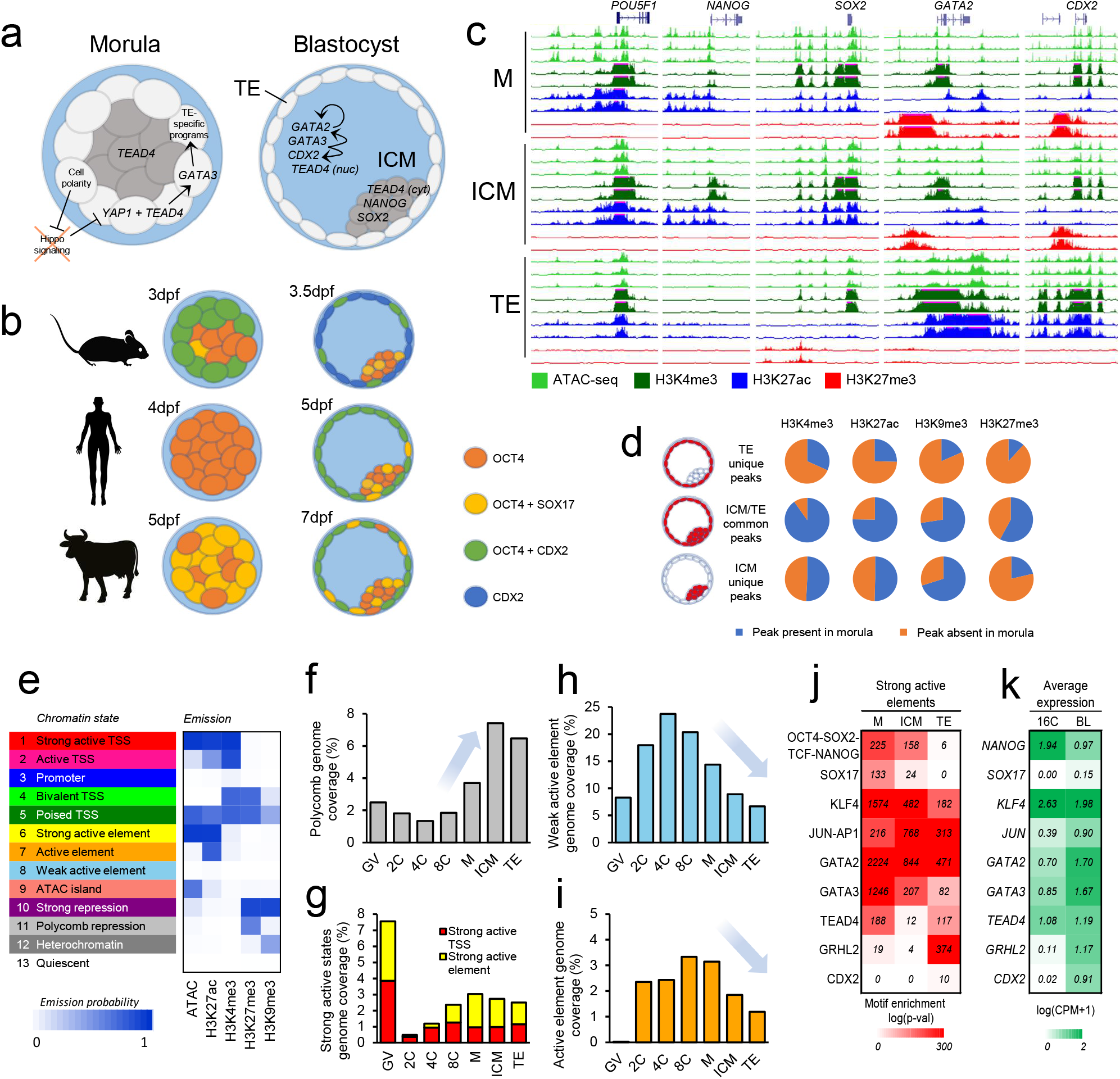
Epigenetic shifts that underscore lineage segregation in the blastocyst. **a**, Conserved mechanisms of TE and ICM specification and maintenance in mammals. **b**, Species-specific differences in TE and ICM markers in morula and blastocysts.The timing of each stage is indicated by days post fertilization (dpf). **c**, Epigenetic profiles of key pluripotency genes and TE-specific markers. Viewing range from 0 to 1.5 CPM. Values exceeding maximum range indicated by pink bars. All biological replicates shown. **d**, For each histone modification, proportion of peaks unique to the ICM, unique to the TE, or shared in common between ICM and TE that were already present in morula. **e**, Chromatin state predictions based on chromatin accessibility and histone modification data. Emission probabilities indicate the likelihood of a given mark occurring in a given state. Genome coverage of **f**, Polycomb repression, **g**, strong active TSS and strong active elements, **h**, weak active elements, and **i**, active elements by developmental stage. **j**, Motif enrichment of selected regulators in strong active elements in M, ICM, and TE. **k**, Average normalized expression of selected regulators in 16-cell embryos (16C) and blastocysts.

We examined our data to identify potential molecular mechanisms responsible for these differences and observed that the *CDX2* locus was Polycomb repressed in morula and ICM, whereas *POU5F1* demonstrated activating signatures in all three cell types (**Fig. 6c**). Expected histone modification profiles were also found at other classical markers of ICM (e.g., NANOG and SOX2) and TE (e.g., GATA2) (**Fig. 6c**). Moreover, the murine-specific TE marker EOMES, which is not detected in human blastocysts, was Polycomb repressed and downregulated in blastocysts, whereas the human-specific TE marker PLAC8, which is not detected in mouse TE, demonstrated stronger H3K27ac in TE and increased expression in blastocysts relative to 16-cell embryos (**Extended Fig. 8a-b**).

Across all of these key loci, both activating and repressive signatures in ICM were very similar to those observed in morula, whereas TE demonstrated a distinct epigenetic profile (**Fig. 6c, Extended Fig. 8a**). The genome-wide distribution of all four histone marks recapitulated this trend, with ICM always falling between morula and TE according to PCA (**Fig. 1a**). The resemblance of morula and ICM was anticipated, because the transition from morula to ICM entails a restriction of pluripotency potential rather than differentiation.

Therefore, we hypothesized that the distinct TE epigenome resulted from *de novo* deposition of histone modifications during lineage specification. Among H3K4me3, H3K27ac, H3K9me3, and H3K27me3 peaks that were TE-specific (e.g., not present in ICM), 68, 75, 82, and 88%, respectively, were not detected in morula (**Fig. 6d, Extended Fig. 8c**). Of note, 79% of ICM-specific H3K27me3 peaks were also newly established during blastocyst formation (**Fig. 6d, Extended Fig. 8c**), suggesting that Polycomb repression plays a major role in specification of both the ICM and TE.

Increased H3K27me3 suggests that an increasingly repressive chromatin landscape is formed as potency declines. This restriction would require reciprocal establishment of enhancer activity to selectively relieve repression and activate cell-specific expression programs. To address whether lineage segregation in the blastocyst follows this model, histone modification and chromatin accessibility datasets were integrated to identify unique chromatin states. The resulting 13 states were annotated based on genome coverage, enrichment at genes and repetitive elements, CpG content and methylation status, and sequence conservation (**Extended Fig. 9**). Overall, chromatin states represented promoter and enhancer activity, constitutive and facultative heterochromatin, bivalency, and quiescence (**Fig. 6e**). As anticipated, repression increased substantially during the morula to blastocyst transition and was largely due to an increase in Polycomb repression (**Fig. 6f**).

Both ICM and TE reached similar levels of repression to that observed in GV oocytes, although oocytes were markedly more enriched for constitutive heterochromatin (H3K9me3), whereas facultative heterochromatin (H3K27me3) was more prevalent in blastocyst lineages (**Extended Fig. 9d**). Concurrently, the active compartment of the genome steadily decreased after EGA, reaching a minimum in TE (**Extended Fig. 9d**). However, the most active regions (e.g., “Strong active elements” and “Strong active TSS”) occupied comparable portions of the genome in morula, ICM, and TE (**Fig. 6g**), whereas regions of intermediate activity (e.g., “Weak active elements” and “Active TSS”) were progressively eliminated, eventually becoming the least prevalent in TE (**Fig. 6h-i**). Overall, these trends suggest that lineage specification in the blastocyst entails an overall refinement of chromatin structure, with targeted repression at cell-specific regions and restriction of active regions.

Despite the comparable genome occupancy of Polycomb repression and highly active elements (e.g., enhancers and promoters) in ICM and TE, the differential distribution of these chromatin states, combined with TF activity, distinguished these two cell lineages. To identify genes specifically repressed or activated in ICM or TE, H3K27me3 and H3K27ac signal were quantified in promoters (2 Kb upstream of TSS) and compared between the two cell types to identify differentially repressed genes (DRG) and differentially activated genes (DAG) (**Extended Fig. 8d**). In TE, DRG (n=1,065 genes) were functionally enriched for calcium ion binding, and DAG (n=1,611 genes) were enriched for cholesterol biosynthesis (**Extended Table 10**). In ICM, DRG (n=949 genes) were enriched for homeobox genes, TFs, and developmental pathways, and DAG (n=1,179 genes) were enriched for TF activity (**Extended Table 10**). Notably, although both DRG and DAG in ICM were enriched for TF activity, different regulators were represented in each gene set (**Extended Table 11**). Moreover, key genes demonstrated both repression and activation in respective cell types. Loci that were DRG in TE and DAG in ICM (n=151 genes) were enriched for cancer pathways and RNA polymerase II core promoter proximal region sequence-specific DNA binding, and loci that were DAG in TE and DRG in ICM (n=309 genes) were enriched for conserved homeoboxes, including CDX2 (**Extended Fig. 8d**).

Considering the overrepresentation of TFs among genes differentially regulated by H3K27me3 and H3K27ac, motif enrichment analysis was conducted on strong active elements – the chromatin state that likely represents active enhancers – to infer key regulators of ICM and TE identity (**Fig. 6j**). Overall, the motifs of ICM- and TE-specific markers demonstrated the expected patterns of enrichment. Key pluripotency factor binding motifs (OCT4-SOX2-TCF-NANOG) were differentially enriched in active ICM enhancers relative to TE. Recognition motifs of SOX17, which is heterogeneously expressed in morula and then restricted to the ICM ^58^, were also enriched in active enhancers in morula and ICM. On the other hand, TEAD4, which is involved in TE-specification in morula ^59^, was specifically enriched in active enhancers in morula and TE, but not in the ICM. These results are consistent with protein localization, as TEAD is markedly cytoplasmic in the ICM and nuclear in TE ^61^. Enrichment of the GRHL2 motif was highly specific to active enhancers in the TE (14% of enhancers contained this motif). Notably, in humans GRHL2 is wide-spread in both morula and TE ^59^, but in bovine there was little evidence of GRHL2 activity at the morula stage, potentially indicating a delay in expression of TE-specific genes in cattle, relative to human. In a similar vein, active enhancers in TE demonstrated very little enrichment for CDX2 motifs, despite notable expression in the blastocyst (**Fig. 6k**) and presence of CDX2 protein ^58,60^. Thus, in terms of genome regulation, CDX2 may play a more prevalent role during later stages of bovine blastocyst development.

In both morula and blastocyst lineages, motifs of KLFs and GATA factors were heavily over-represented in active enhancers. However, because of similarities in binding motifs between family members, it remains unclear which specific members are active in a given cell type. For example, whereas GATA3 induces TE fate ^59,62,63^, GATA6 and GATA4 are implicated in primitive endoderm specification and maintenance ^64–66^. Among KLFs, which generally play roles in pluripotency maintenance, KLF2 marks murine epiblast but is absent in humans, whereas KLF17 is widespread in human blastocysts but absent in mouse ^67–69^. The enrichment of KLF motifs in bovine morula, ICM, and TE is reminiscent of the widespread expression KLF17 in human embryos, but it remains to be determined which member(s) of the KLF family are active in bovine; several candidates are expressed at the blastocyst stage, including *KLF6*, which undergoes a 14-fold increase in expression between the 16-cell and blastocyst stages (**Extended Fig. 8e**).

Similar to KLF and GATA factors, the motif of JUN-AP1, a factor which was recently implicated as a gatekeeper to reprogramming of pluripotency ^70^, was also enriched in active enhancers in morula, ICM, and TE. However, based on TF footprinting analysis of chromatin accessibility data, which measures TF activity based on reduced DNA cleavage at motifs actively bound by TFs, the activity of JUN and FOS factors – which together form the AP-1 complex – was markedly increased in ICM relative to morula (**Extended Fig. 8e**) and was accompanied by an increase in expression of both JUN and FOS factors (**Extended Fig. 8g**). Moreover, the morula to ICM transition entailed a loss in homeobox activity (**Extended Fig. 8e**). These shifts in TF activity distinguished these two pluripotent cell types, which otherwise demonstrated strong epigenetic similarities, both in terms of histone modification profiles (**Fig. 6a**) and motif enrichment in active enhancers (**Fig. 6j**).

Overall, we find that lineage specification entails further refinement of the epigenome, including increased Polycomb repression at specific loci, especially those related to future developmental pathways, and resolution of regions with intermediate or weak activity. Because the transition from pluripotency to a differentiated state is generally irreversible, it is possible that by refining activity to certain genomic regions, the potential to activate others is consequently lost. Finally, these epigenetic shifts were complimented by differential expression and activity of key TFs, including known markers of human ICM and TE, which further supports the use of bovine embryos as a relevant model for human development.

## Conclusion

Accumulating evidence indicates that the mechanisms underlying the MZT and blastocyst formation in rodents are distinct from those employed by other mammals. Such differences highlight the need to establish additional animal models, such as bovine, to investigate the molecular basis for preimplantation development in humans. Although certain phenomena that are present in both ungulates and rodents (e.g., ncH3K4me3) are uniquely absent in humans ^20^, cattle and human embryos demonstrate striking similarities in developmental timing, morphology, and regulators of both EGA and blastocyst formation^12,58,60,71^. For example, bovine embryos may be an appropriate model to study the function of genes not expressed in mouse, such as PLAC8, for which the function in TE remains unknown. To this end, the atlas of chromatin states produced by this study will be an invaluable resource for future research in mammalian embryos, ranging from functional studies of specific genes, to broadening our understanding of genome reprogramming during the preimplantation period.

## Acknowledgements

Funding for this project was provided by NIH grant HD070044 to PJR and RMS. MMH and ABG were funded by the REVIVE Labex (Investissement d’Avenir, ANR-10-LABX-73) and supported by the PHASE Department of the French National Research Institute for Agriculture, Food and Environment (INRAE). The authors declare no competing financial interests.

## Data availability

The following published datasets were accessed through the NCBI Gene Expression Omnibus (GEO) repository: RNA-seq data for bovine oocytes and embryos (accession no. GSE52415 ^72^ and GSE110040 ^73^), ATAC-seq data for bovine oocytes and embryos (GSE143658 ^12^), CUT&Tag data for bovine fibroblasts (GSE171104), and CUT&RUN data for bovine oocytes and blastocysts (GSE163620 ^20^). CUT&Tag and ATAC-seq data produced in this study have been deposited in the NCBI GEO database (GSE193640). A UCSC track hub is available to view predicted chromatin states, ATAC-seq, and CUT&Tag read depth (https://genome-euro.ucsc.edu/s/mmhalstead/Bovine_Embryo_Epigenome).

## Author contributions

CZ, RMS, and PJR designed the experiments. CZ conducted the experiments. MH, CZ, ABG, RMS, and PJR performed data analysis and wrote the manuscript. All authors read and approved of the final manuscript.

## Materials and Methods

### Collection of bovine oocytes and preimplantation embryos

Bovine ovaries were collected from a local slaughterhouse and transported to the laboratory in 0.9% NaCl solution, complying with the guidelines of University of California Davis relevant ethical regulation for animal research. Cumulus-oocyte complexes (COCs) were aspirated from antral follicles (2-10 mm in diameter) with a 10 mL syringe and then washed in M199 (Sigma, M7653) containing 2% (v/v) fetal bovine serum (FBS, Hyclone). Only COCs with at least three layers of compact cumulus cells and evenly granulated cytoplasm were selected for maturation *in vitro*. After washing three times in M199 supplemented with 2% (v/v) FBS, COCs were cultured for 22-24 h in BO-IVM media (IVF Bioscience, 71001) in an atmosphere of 5% CO_2_ in air at 38.5 °C. After maturation *in vitro*, MII oocytes were washed three times in SOF-IVF medium (107.7 mM NaCl, 25.07 mM NaHCO_3_, 7.16 mM KCl, 1.19 mM KH_2_PO_4_, 1.17 mM CaCl_2_, 0.49 mM MgCl_2_, 5.3 mM sodium lactate, 0.20 mM sodium pyruvate, 10 μg/ml heparin, 0.5 mM fructose, 1x nonessential amino acids, 6 mg/ml BSA, 5 μg/ml gentamicin), and then transferred to 90 μL drops (50 COCs per drop) of SOF-IVF covered with mineral oil (Vitrolife, 10029). Frozen semen was thawed in water at 37 °C for 1 min. Sperm were selected using density gradient centrifugation, washed once using TALP-Sperm medium (100 mM NaCl, 25 mM NaH_2_CO_3_, 3.1 mM KCl, 2.1 mM CaCl_2_, 0.29 mM NaH_2_PO_4_, 0.4 mM MgCl_2_, 21.6 mM sodium lactate, 10 mM Hepes, 6 mg/ml BSA, 5 μg/ml gentamicin), counted, and 10 μL of 2×10^6^ sperm/mL were added to each drop containing matured oocytes. After incubating for 6 h in an atmosphere of 5% CO_2_ in air at 38.5 °C, cumulus cells were removed by vortexing for 5 min and zygotes were washed three times in BO-IVC media (IVF Bioscience, 71005) and then transferred to 100 μL drops (50 zygotes per drop) of BO-IVC media covered with mineral oil. For each collection, 50 zygotes were set aside and cultured to the blastocyst stage and only collections for which the incidence of blastocyst formation was at least 20% were used.

Cumulus cells were removed from COCs by vortexing for 5 min and GV oocytes were collected for CUT&Tag library construction. Embryos at the 2-, 4-, 8-cell (cultured in the presence or absence of 50 μg/mL a-amanitin (Sigma, A2263)), morula, and blastocyst stages were collected at 32, 44, 56, 122, and 172 h post-insemination, respectively. For ICM collection, blastocysts were collected at day 7 and subjected to immunosurgery with anti-bovine serum antibody (Sigma, B8270) and guinea pig complement serum (Innovative Research, IGP-COMPL-21249) as previously described ^12^. For TE separation, blastocysts were collected at day 7 and subjected to surgery under a microscope, cutting with a micro scalpel (Feather, 72046-30). GV oocytes and embryos were treated with 0.5% pronase to remove the zona pellucida, and then used for library construction. Approximately 500 oocytes or 1,000-1,500 blastomeres were collected for each CUT&Tag or ATAC-seq library.

### CUT&Tag library construction and sequencing

CUT&Tag was performed following the manufacturer’s instructions (Hyperactive In-Situ ChIP Library Prep Kit, TD901, Vazyme) with modifications. In brief, bovine oocytes and embryos were incubated with concanavalin-coated magnetic beads for 20 min on a thermomixer at 400 rpm at room temperature (RT). The samples then were incubated with a primary antibody (1:50 dilution of a rabbit polyclonal anti-H3K4me3 (Cat# C15410003), H3K27ac (Cat# C15410196), H3K27me3 (Cat# C15410195), or H3K9me3 (Cat# C15410193) from Diagenode) overnight at 4 °C on a nutator, then with the secondary antibody (1:100 dilution, ABIN101961, Antibodies online) for 1 h on a nutator at room temperature (RT), then with hypoactive pA-Tn5 transposon for 1 h on a nutator at RT. To perform targeted digestion, 300 µL tagmentation buffer was then added, and samples were incubated at 37 °C for 1 h. The reaction was terminated by adding 10 µL of 0.5 M EDTA, 3 µL of 10% SDS and 2.5 µL of 20 mg/mL proteinase K to each tube, which were then incubated at 50°C for 1 h. DNA extraction was performed by adding 300 µL of PCI (Phenol: Chloroform: Isoamyl alcohol = 25:24:1) to each tube and mixed by full-speed vortexing, after which the sample was transferred to a phase-lock tube (1038987, Qiagen) and centrifuged for 3 min at RT at 16,000 x g. Then, 300 µL chloroform was added followed by centrifugation for 3 min at RT at 16,000 x g. The aqueous phase was removed and transferred to a 1.5 mL tube to which 750 µL of 100% ethanol was added, the tube inverted 10 times, and then centrifuged for 30 min at 4 °C at 16,000 x g. The supernatant was removed and 1 mL of 100% ethanol added, followed by centrifugation for 5 min at 4 °C at 16,000 x g. The supernatant was removed and the tube then allowed to dry in air. TE buffer (20 µL) was added and PCR was performed using NEBNext High-Fidelity 2X PCR Master mix (New England Biolabs, M0541) as follows: 58 °C for 5 min, 72 °C for 5 min, 98 °C for 45 s, followed by 14 cycles of 98 °C for 15 s and 60 °C for 10 s, with a final extension at 72 °C for 1 min. Purification of PCR products was performed using Ampure XP beads (Bechman Coulter, A63881) and libraries were pooled for sequencing on the Illumina NextSeq platform to generate 40 bp paired-end reads.

### ATAC-seq library construction and sequencing

Pools of zona-free blastocysts, ICM and TE were collected for ATAC-seq ^23^ library construction from three separate collections, respectively, and transferred to cold lysis buffer (10 mM Tris-HCl pH7.4, 10 mM NaCl, 3 mM MgCl_2_, and 0.1% IGEPAL CA-630). The samples were incubated on ice for 5 min, and then centrifuged using a swinging bucket rotor for 10 min at 500 x g at 4 °C. The supernatant was removed and the pellet was resuspended in 50 μl of transposition reaction mix (25 μl 2x TD buffer (Nextera DNA Library Prep Kit, Illumina), 2.5 μl TDE1 enzyme (Nextera DNA Library Prep Kit, Illumina), 22.5 μl ddH_2_O) by pipetting up and down three times. Transposition reactions were incubated at 37 °C for 60 min in a thermomixer at 300 rpm. Reaction products were purified with a MinElute PCR purification kit (Qiagen, Germany) and eluted in 20 μl EB buffer. Transposed DNA was amplified with NEBNext High-Fidelity 2X PCR Master mix (New England Biolabs, M0541) as follows: 58 °C for 5 min, 72 °C for 5 min, 98 °C for 45 s, followed by 12 cycles of 98 °C for 15 s and 60 °C for 10 s, with a final extension at 72 °C for 1 min. Libraries were purified with Ampure XP beads (Bechman Coulter, A63881) and were pooled for sequencing on the Illumina NextSeq platform to generate 40 bp paired-end reads.

### CUT&Tag and ATAC-seq data processing

Raw reads were trimmed with Trim Galore (v0.4.0) and Cutadapt (v1.8.3) ^74^ with options “-q 20 -stringency 1 -length 10” to remove adaptor sequences and low-quality reads (q < 20). Trimmed reads were then mapped to the cattle reference genome (ARS-UCD1.2) using BWA (v0.7.9a) ^75^ in ‘mem’ mode and with default parameters. Duplicate reads were removed using PicardTools (v2.26.10) and low-quality mapped reads (q < 15) were removed using SAMtools (v1.9) ^76^ to obtain the final informative reads which were used for downstream analyses.

### RNA-seq data processing

RNA-seq data of bovine oocytes and embryos were obtained from two previous studies ^72,73^. Raw reads were trimmed with Trimmomatic (v0.33) ^77^ to remove low-quality leading and trailing bases (3 bases) and adapter sequences, allowing for 2 seed mismatches, a palindrome clip threshold of 30, and simple clip threshold of 10. Trimmed reads were mapped to the cattle reference genome (ARS-UCD1.2) with STAR (v2.7.2a) ^78^ with default parameters, except ‘–seedSearchStartLmax 30 –outFilterScoreMinOverLread 0.85’. Low-quality alignments (q < 5) were filtered using SAMtools. HTseq-count (v0.10.0) ^79^ was used to calculate the raw count for genes in the Ensembl (v104) annotation with parameters ‘— mode=intersection-nonempty –type=exon.’ Differentially expressed genes (DEG) were identified using the DESeq2 R package (v1.34.0) ^80^. To identify maternal and embryonic gene sets, DEG (log2FC>1 and FDR<0.01) were determined from pair-wise comparisons of MII oocytes, 8-to-16-cell embryos, and 8-to-16-cell embryos cultured in the presence of a-amanitin, as described previously ^73^. To identify minor EGA genes, DEG (log2FC>1 and FDR<0.05) were determined between MII oocytes and 4-cell embryos ^72^. A looser FDR cutoff was used to capture subtler differences in gene expression. Counts were normalized by counts per million (CPM) and z-score transformed for visualization.

### DNA methylation data processing and identification of PMDs

DNA methylation data were obtained from a previous study ^32^. Raw reads were aligned to the cattle reference genome (ARS-UCD1.2) using Bismark (v0.14.3) ^81^, with options “–bowtie2 – pbat.” Context-dependent CpG methylation was then extracted using the “bismark_methylation_extractor” command with options “–s –bedGraph –cytosine_report.” To identify PMDs, the genome was first binned into 1 Kb windows with the BEDtools (v2.27.1) ^82^ command makewindows. Bins that contained 5 or more CpGs, and which had an average methylation level less than or equal to 0.5 in GV oocytes, were considered hypomethylated windows. These were merged, allowing for a maximum gap between windows of 10 Kb. Regions that fell within 2.5 Kb of TSS were excluded from the final set of PMDs using BEDtools subtract. Methylation status of genomic features (e.g., gene bodies, peaks, chromatin states, etc.) was determined by taking the average methylation level of all CpGs that overlapped the given feature.

### Reproducibility of CUT&Tag and ATAC-seq data

Informative reads were converted to bigwig format and signal was normalized by counts per million (CPM) with bamCoverage from the DeepTools suite (v3.4.3) ^83^ with default parameters. Resulting bigwig files were uploaded to the UCSC genome browser ^84^ for track visualization. The viewing range was set from 0 to 1.5 CPM, and values exceeding the maximum range were indicated by pink bars. The smoothing window was set to 4 pixels, and windowing function was set to “mean + whiskers”, which shows the average signal in the darkest shade, one standard deviation away from the mean in a medium shade, and the maximum and minimum in the lightest shade. To calculate the correlation between libraries and conduct principal components analyses, bigwig files were first consolidated using the multiBigWigSummary function from Deeptools, using a bin size of 500bp for activating marks (ATAC-seq, H3K4me3, H3K27ac) and 10 Kb for repressive marks (H3K9me3, H3K27me3). The output was then piped to plotPCA with parameters ‘-transpose -log2 -ntop 100000’ (active marks) or ‘-transpose -log2 -ntop 50000’ (repressive marks), and plotCorrelation with parameters ‘-corMethod pearson -skipZeros’. Finally, files containing normalized signal (RPKM; reads per kilobase million) of published CUT&RUN H3K27me3 data for bovine GV oocytes and blastocysts were downloaded from GSE163620 ^20^. Using the liftOver tool from the UCSC tool suite, these signal files were converted to the most recent genome assembly. To compare these data to CUT&Tag, the H3K27me3 libraries from GV oocytes and blastocysts were also normalized by RPKM using bamCoverage from DeepTools, and the multiBigWigSummary and plotCorrelation functions were implemented as previously described to determine correlation between libraries.

### Peak calling

To identify regions with signal enrichment, or “peaks,” for each library, we first compared two different methods for peak calling: Epic2 ^85^ and Macs2 ^86^, using different parameters for narrow and broad epigenetic marks (**Extended Table 12**). Peaks from biological replicates were compared with the BEDtools jaccard command. We found that Epic2 generated more consistent peak calls between biological replicates, was less sensitive to differences in read depth, and more logically captured regions of enrichment, especially for repressive marks (**Extended Fig. 10**). Therefore, Epic2 was selected as the optimal peak caller, using different parameters for narrow and broad peaks (**Extended Table 12**). Unless otherwise stated, only the overlapping regions, identified by BEDtools intersect, of peaks called in both biological replicates were identified as ‘true’ peaks and used for further analyses. Peaks were classified as genic if they overlapped promoters (2 Kb regions upstream of TSS) or gene bodies. Otherwise, peaks were considered intergenic. Genomic coverage of peaks was calculated using BEDtools genomecov.

### Enrichment of CUT&Tag and ATAC-seq signal

To visualize the signal intensity of histone modifications and chromatin accessibility at specific genomic regions (e.g., TSS, gene bodies, peaks, PMDs), normalized signal from individual libraries or pooled replicates was analyzed using the computeMatrix function of DeepTools with option ‘--skipZeros’. Signal at PMDs was visualized using a bin size of 100bp. Signal at genes and peaks was visualized using a bin size of 10bp.The output was then piped to the plotProfile and plotHeatmap functions from DeepTools. In cases where regions were clustered, the option “—kmeans” was used with plotHeatmap and cluster members were extracted using “—outFileSortedRegions.” Signal at PMDs was summarized by taking the average of all 100bp bins for a given PMD, and then calculating the mean signal and standard error of all PMDs in a given cluster for a given developmental stage.

### Comparison of DNA methylation and intragenic H3K27me3 signal

For H3K27me3 libraries, signal in gene bodies was quantified using HTseq-count with parameters “—mode=intersection-nonempty –type=gene.” Raw counts were processed using DESeq2 to obtain log2FC values between GV oocytes and 2-cell embryos. Correlations between gene body H3K27me3 (log2FC) and change in CpG methylation (2-cell versus GV oocytes) were calculated by linear regression.

### Motif enrichment analysis

Intergenic regions were used for motif enrichment analysis with the findMotifsGenome.pl script in HOMER (v4.11.1) ^87^ using parameters ‘-size given -mask’. Known motifs were reported based on p-value.

### Functional enrichment analysis

Gene sets were identified based on overlap with peaks or chromatin states using the intersect function from BEDtools. Functional enrichment analysis of gene sets was performed using the Database for Annotation, Visualization and Integrated Discovery (DAVID, v2021q4) ^88,89^. Terms with an FDR less than 0.05 were considered significant.

### Identification of differentially activated and repressed genes in ICM and TE

For H3K27me3 and H3K27ac libraries, signal in promoters (2 Kb regions upstream of genes) was quantified using HTseq-count in mode “intersection-nonempty.” Genes with differential promoter signal (log2FC>1 and FDR<0.05) between ICM and TE were identified using DESeq2, and counts were z-score transformed for visualization.

### Transcription factor footprinting analysis

Transcription factor footprints were identified from ATAC-seq data using HINT from the Regulatory Genomic Toolbox ^90^ as previously described^12^. For footprint detection, alignments from biological replicates were pooled using the merge function from SAMtools, and peaks were called from pooled alignments using Epic2. TFs with a significant difference in binding activity between two cells types were reported (two-tailed *t* test; p < 0.05).

### Chromatin state annotation

ChromHMM (v1.22) ^91^ was used to train a chromatin state model incorporating CUT&Tag (H3K4me3, H3K27ac, H3K27me3, and H3K9me3) and ATAC-seq from all developmental stages except for blastocyst. The biological replicates of the same developmental stage were merged into one file using the merge function from SAMtools. Specifically, the alignment files from all developmental stages were first binarized using the ‘BinarizeBam’ command with the default parameters. The output was then piped to generate the segmentation model using the ‘LearnModel’ command with the default parameters. Multi-state models were trained and the 13-state model was finally selected as it exhibited the maximum number of states with distinct chromatin mark combinations. Fold enrichment of chromatin states at genomic features was calculated using the ‘OverlapEnrichment’ command. Fold enrichment was calculated as (C/A)/(B/D), where A represented the bases in a chromatin state, B the bases in a genomic feature, C the number of bases overlapped between a chromatin state and genomic feature, and D the bases in the genome. Chromatin state fold enrichment was calculated for gene elements (Ensembl v104 annotation), repetitive elements (RepeatMasker downloaded from UCSC genome annotation database), and mammalian conserved elements identified from multiple sequence alignments using the Genomic Evolutionary Rate Profile (GERP) software based on 103 mammals (ftp://ftp.ensembl.org/pub/release-100/bed/ensembl-compara/103_mammals.gerp_constrained_element/).

---

**Extended Figure 1.**
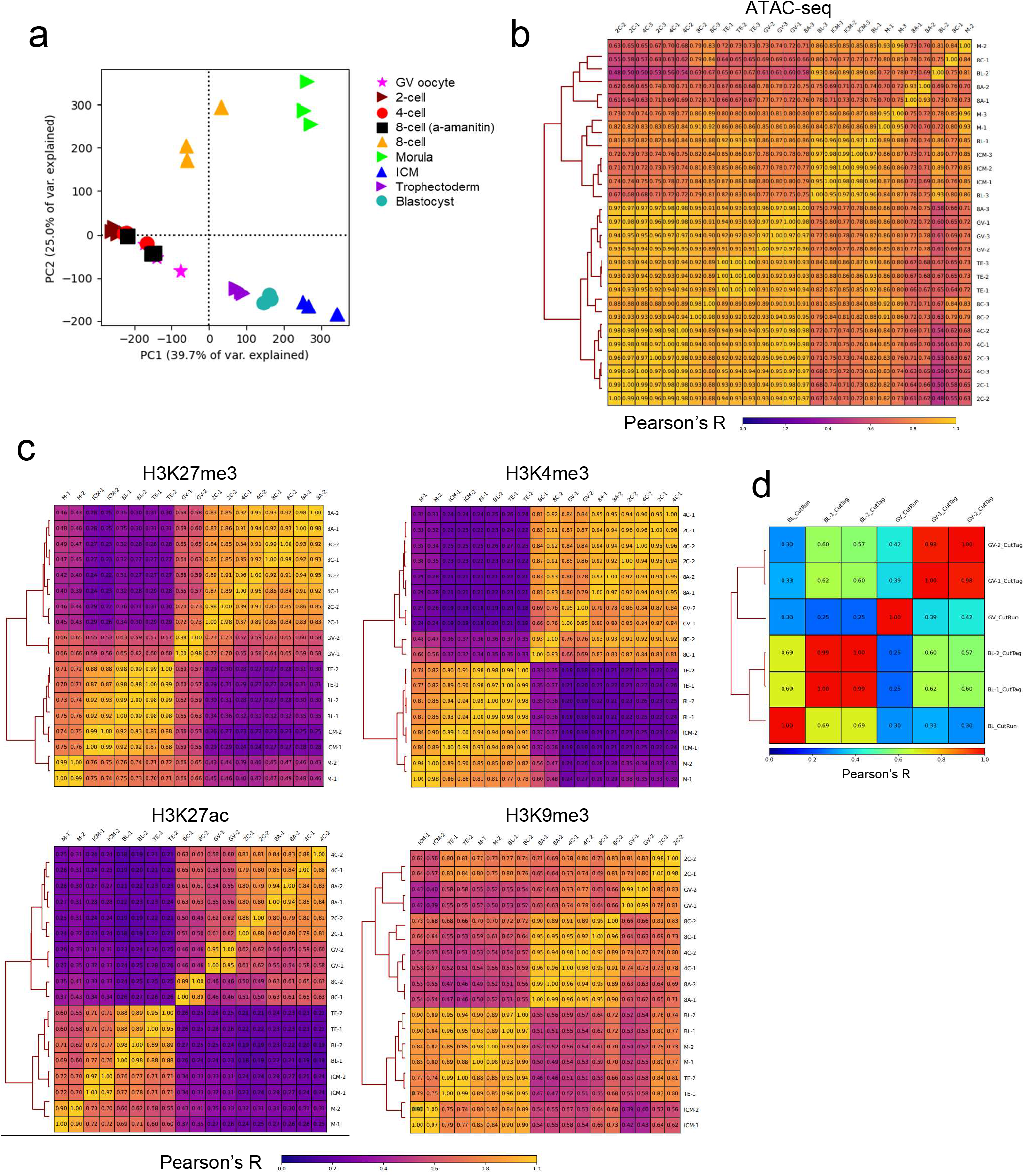
Quality control of CUT&Tag and ATAC-seq libraries. **a**, Principal components analysis (PCA), comparing ATAC-seq libraries (BL, ICM, TE) with published data (GV, 2C, 4C, 8C, 8A, M). **b**, Pearson correlation coefficient (Pearson’s R) of ATAC-seq libraries, based on genome-wide normalized signal (CPM) in 500 bp windows. **c**, Correlation of CUT&Tag libraries based on CPM in either 500 bp windows for narrow marks (H3K4me3 and H3K27ac) or 10Kb windows for broad marks (H3K27me3 and H3K9me3). **d**, Comparison of published CUT&RUN H3K27me3 profiles of bovine GV oocytes and blastocysts with CUT&Tag libraries. Pearson correlation coefficient between libraries based on genome-wide normalized signal (reads per kilobase million; RPKM) in 10Kb windows.

**Extended Figure 2.**
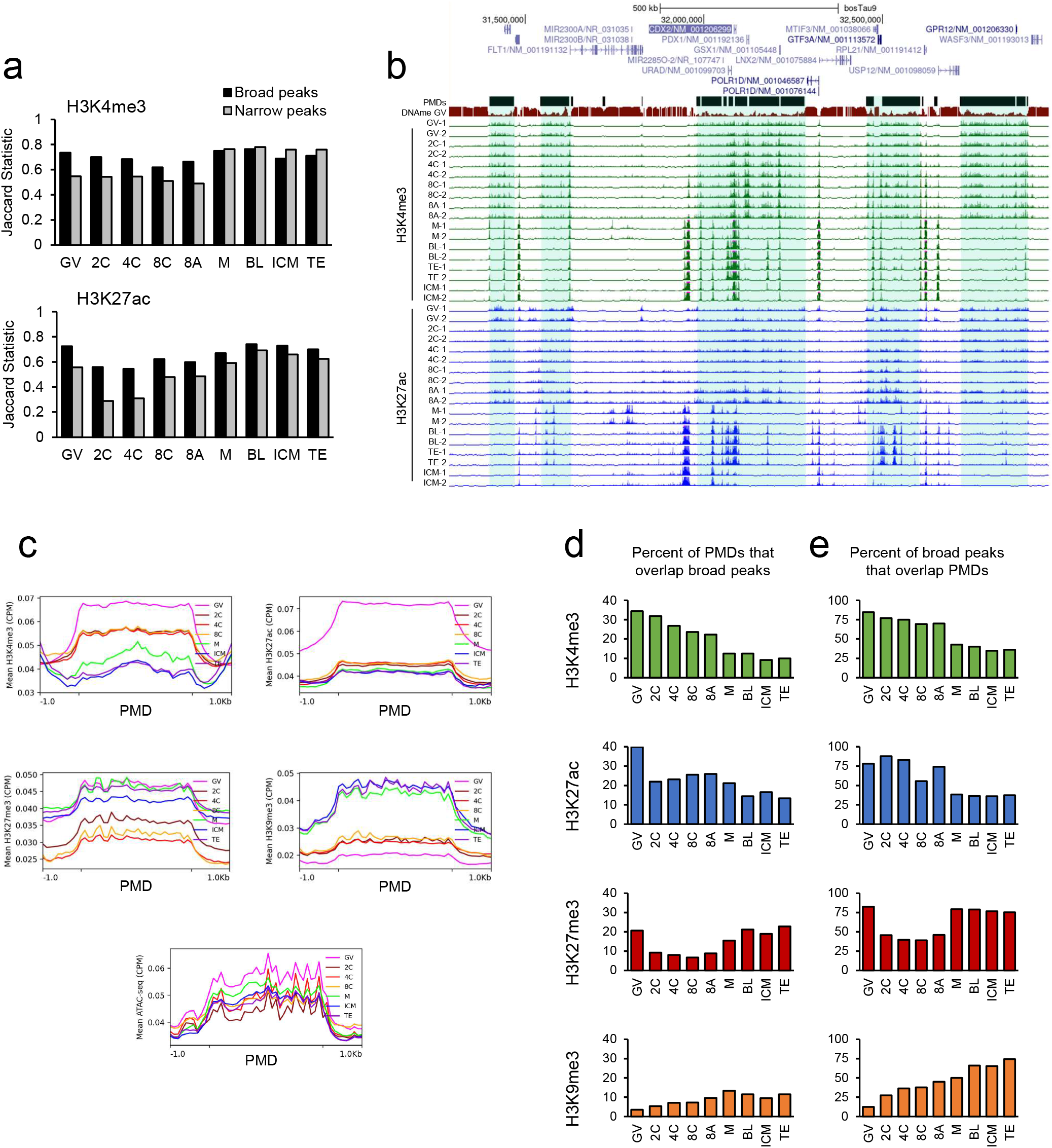
Identification of noncanonical broad H3K4me3 and H3K27ac relative to partially methylated domains (PMDs). **a**, Comparison of peak sets from biological replicates identified using either ‘broad’ or ‘narrow’ peak calling parameters. Jaccard statistic measures base-pair overlap between two peak sets, ranging from 0 (no overlap) to 1 (complete overlap). **b**, Gene track image centered on the *CDX2* locus, with representative PMDs predicted from GV oocyte CpG methylation data (DNAme). Highlighted regions correspond to PMDs, and overlap with broad H3K4me3 and H3K27ac domains. Viewing range from 0 to 1.5 CPM. Values exceeding maximum range indicated by pink bars. **c**, Average normalized signal (CPM) at PMDs for each histone modification and ATAC-seq. PMDs scaled to 3Kb ± 1Kb up- and downstream. **d**, Percent of PMDs overlapped (by at least 1bp) by broad peaks of a given histone modification. **e**, Percent of broad peaks for a given histone modification overlapped (by at least 1bp) by PMDs.

**Extended Figure 3.**
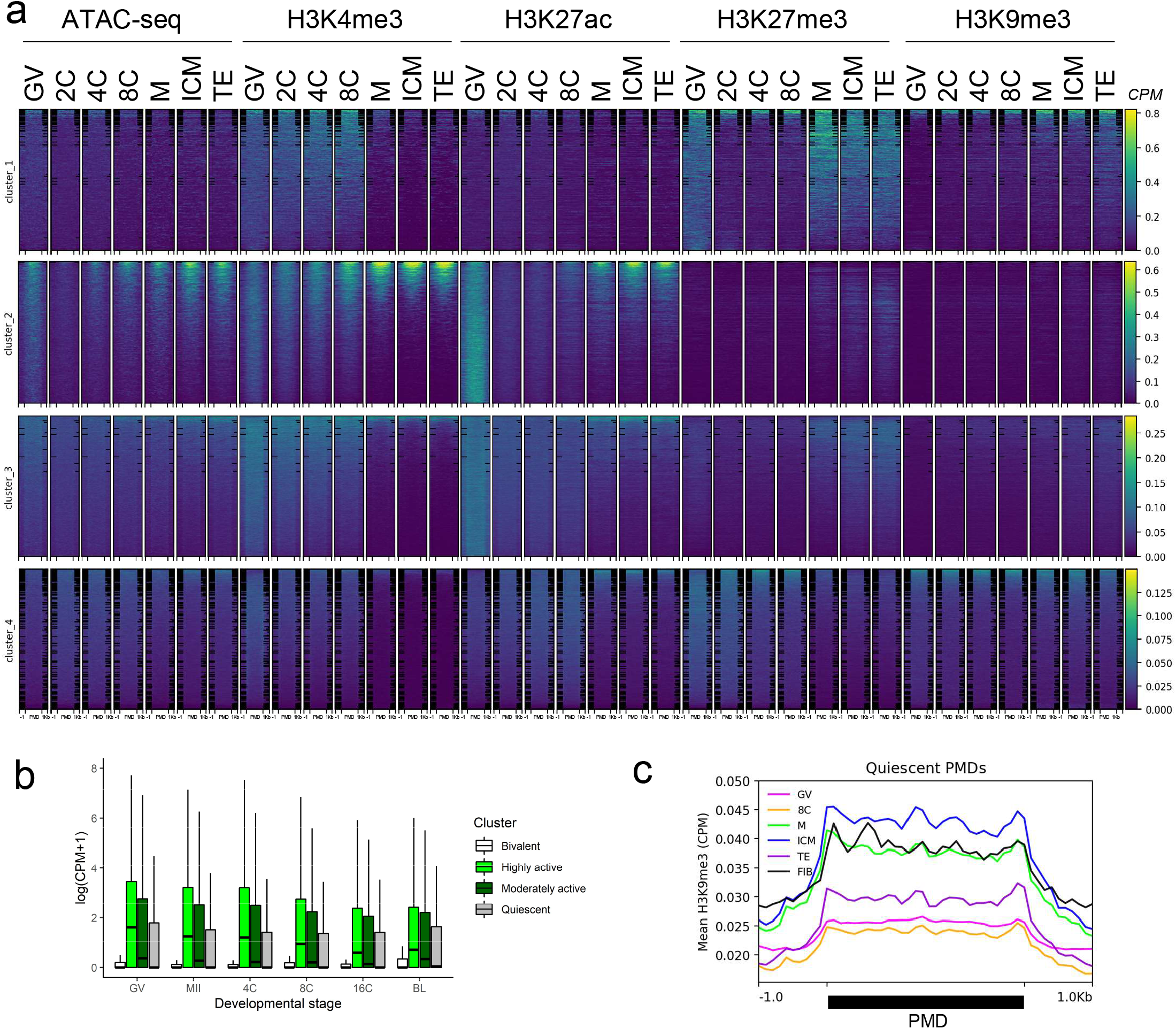
Epigenetic signature of PMD clusters identified in GV oocytes. **a**, Normalized signal (CPM) of each epigenetic mark is shown for GV oocytes (GV), 2-cell (2C), 4-cell (4C), 8-cell (8C), morula (M), inner cell mass (ICM), and trophectoderm (TE) in PMDs. PMDs scaled to 3Kb, with 1Kb up- and downstream regions shown. **b**, Normalized expression (CPM) of genes marked by each PMD cluster in GV, MII oocytes, 4C, 8C, 16-cell embryos (16C) and BL. Boxplots indicate the median and interquartile range (IQR), and whiskers span 1.5 times the IQR. Outliers not shown. **c**, Average normalized H3K9me3 signal (CPM) at quiescent PMDs in oocytes, embryos, and fibroblasts (FIB), scaled to 3Kb, and showing 1Kb up- and downstream.

**Extended Figure 4.**
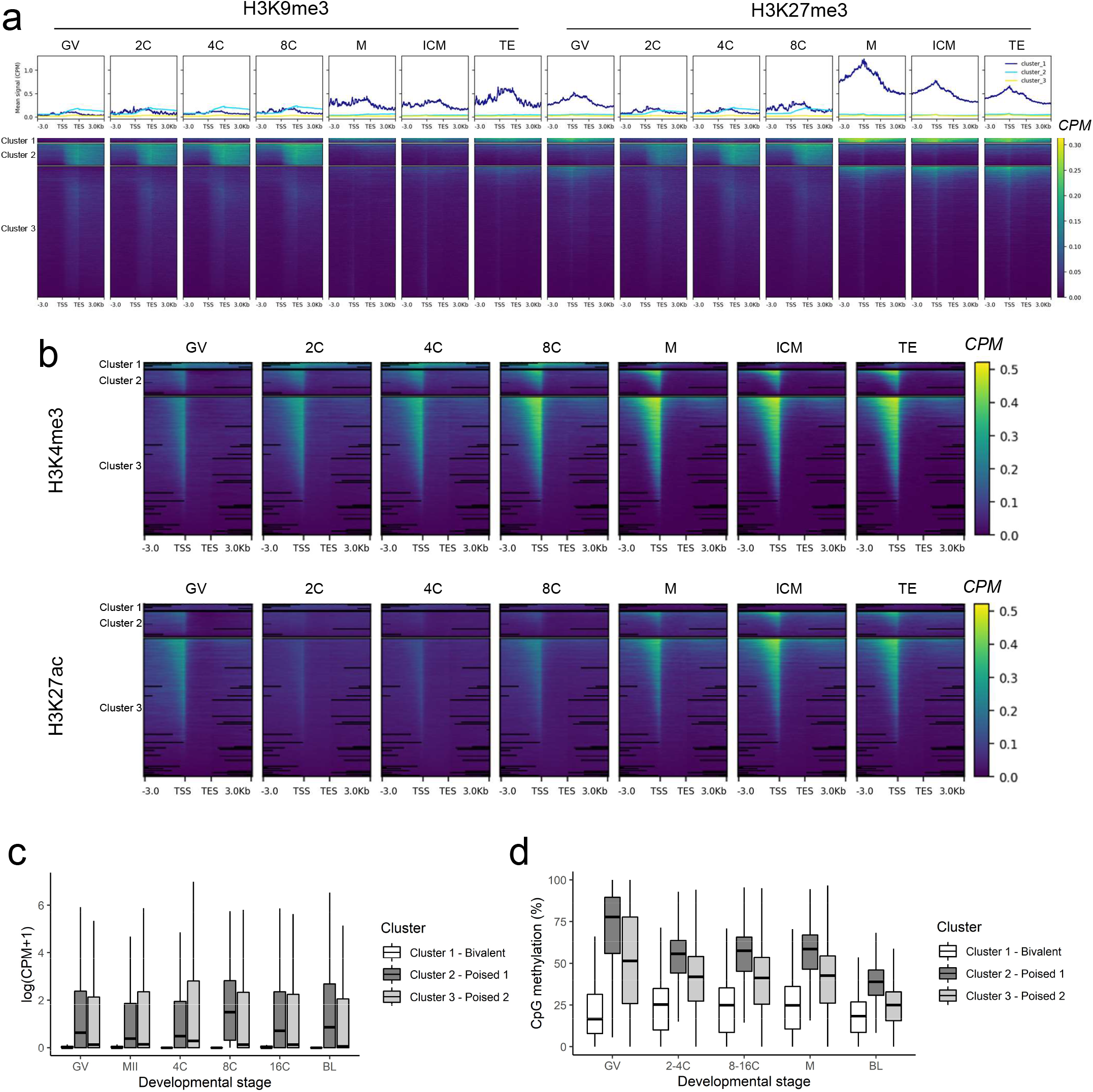
Clustering of transcripts (k=3) based on normalized H3K27me3 and H3K9me3 signal (CPM). **a**, Clustering results based only on repressive marks. **b**, signal of H3K4me3 and H3K27ac at the three clusters defined in (a). **c**, Normalized expression (CPM) of genes marked by each cluster throughout development. **d**, Changes to gene body methylation (CpG) of genes marked by each cluster throughout development. Boxplots indicate the median and interquartile range (IQR), and whiskers span 1.5 times the IQR. Outliers not shown.

**Extended Figure 5.**
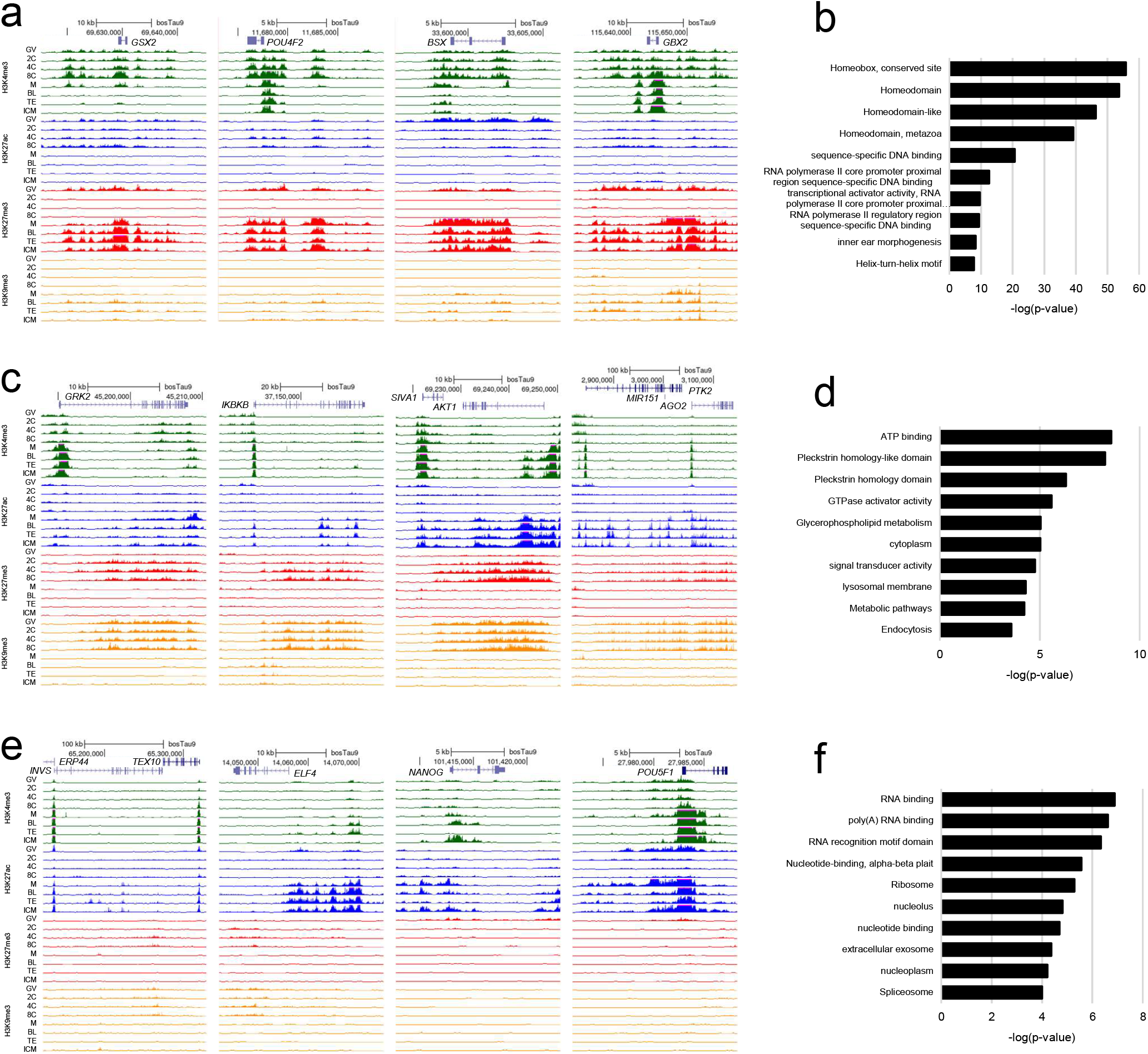
Examples and functional enrichment of transcripts belonging to each cluster defined by H3K27me3 and H3K9me3 signatures. **a**, Example loci belonging to bivalent transcripts (cluster_1). **b**, Functional enrichment of transcripts belonging to bivalent transcripts (cluster_1). **c**, Example transcripts belonging to poised-1 transcripts (cluster_2). **d**, Functional enrichment of transcripts belonging to poised-1 transcripts (cluster_2). **e**, Example transcripts belonging to poised-2 transcripts (cluster_3). **f**, Functional enrichment of transcripts belonging to poised-2 transcripts (cluster_3). For gene tracks, viewing range limited from 0 to 1.5 CPM. Values exceeding maximum range indicated by pink bars. One replicate shown per stage and mark. For functional enrichment, top 10 Gene Ontology terms, INTERPRO terms, and KEGG pathways, based on FDR, are shown.

**Extended Figure 6.**
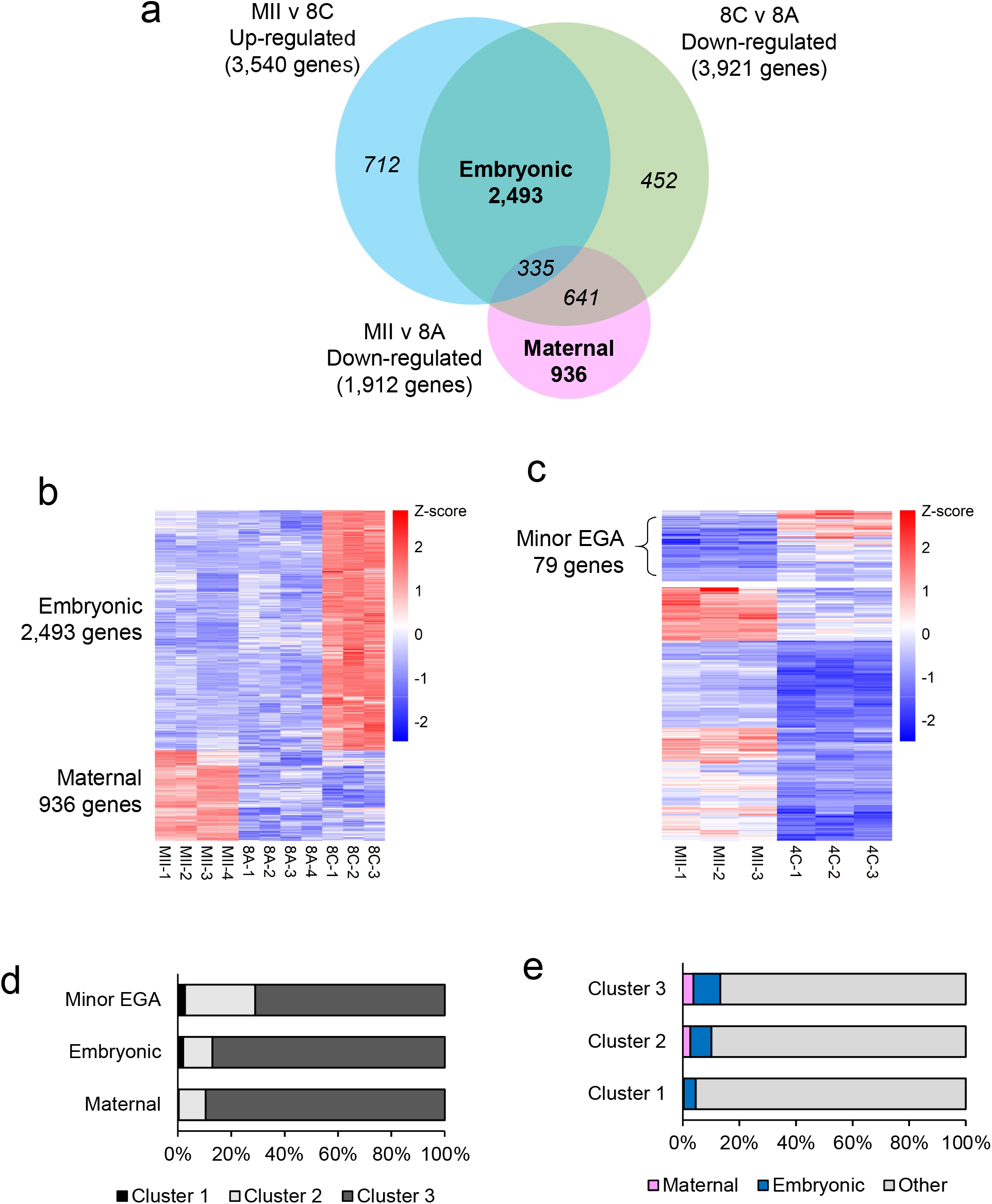
Identification of maternal and embryonic gene sets by differential expression (DE) analysis, and comparison with transcript clusters based on intragenic H3K9me3 and H3K27me3 signal. **a**, Identification of maternal and embryonic gene sets based on comparison of DE genes (log2FC > 1, FDR < 0.01) between MII oocytes (MII), 8-cell embryos (8C), and 8-cell embryos treated with a-amanitin (8A). **b**, Z-scores of embryonic and maternal gene sets. **c**, Z-scores of minor EGA genes identified based on DE between MII oocytes and 4-cell embryos (log2FC > 1, FDR < 0.05). A looser FDR cut-off was used to capture subtler differences in gene expression. **d**, Percent of each gene set (maternal, embryonic, or minor-EGA) that belonged to each transcript cluster, defined by the abundance of H3K27me3 and H3K9me3 (Extended Fig. 4). **e**, Percent of each transcript cluster that belonged to the maternal or embryonic gene sets.

**Extended Figure 7.**
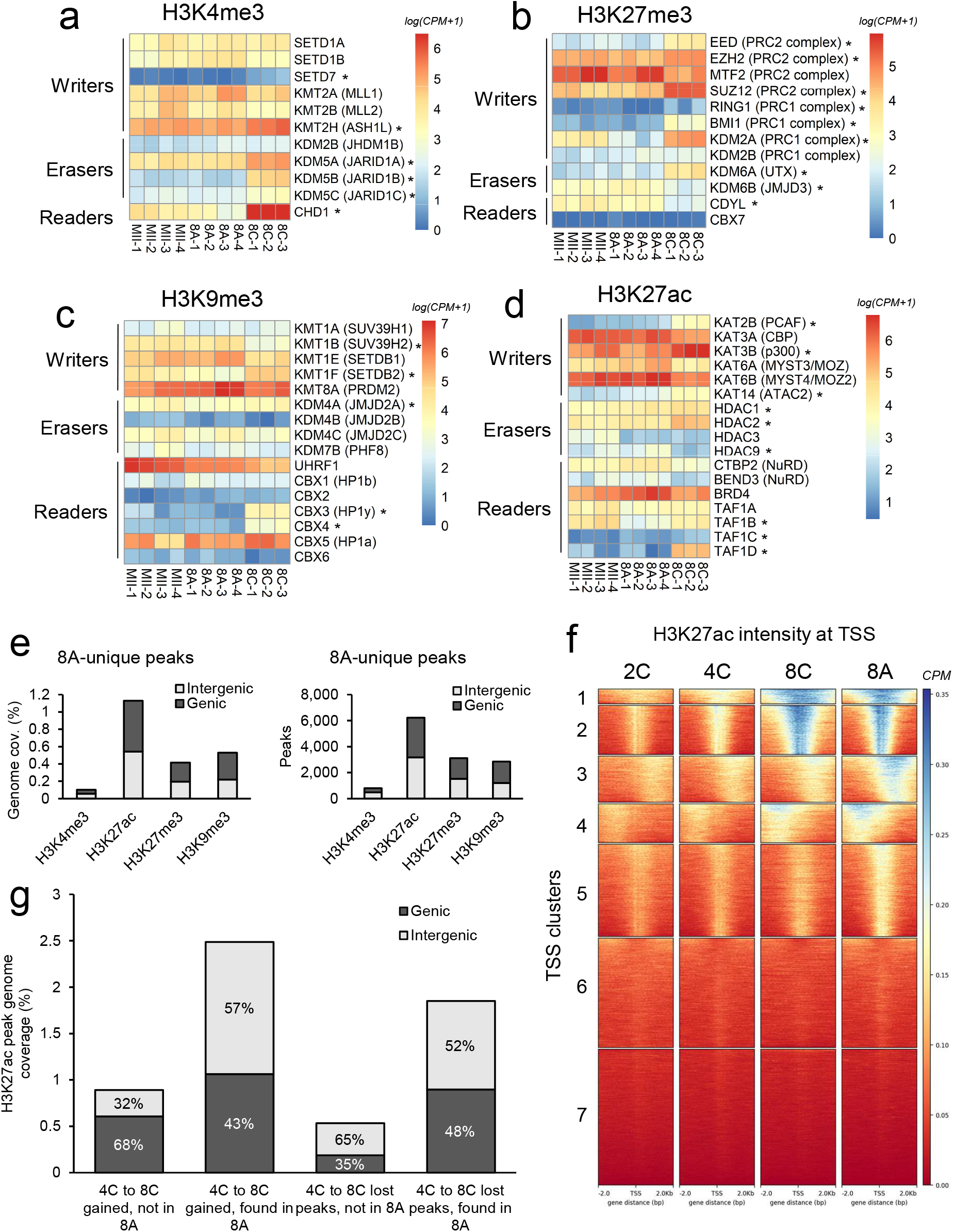
Epigenetic and transcriptomic perturbations in transcription-inhibited 8-cell embryos. Normalized expression (CPM) of writers, readers, and erasers for **a**, H3K4me3, **b**, H3K27me3, **c**, H3K9me3, and **d**, H3K27ac in MII oocytes, 8-to-16-cell embryos cultured in the presence of a-amanitin (8A), and control 8-to-16-cell embryos (8C). Genes marked “*” were differentially expressed between 8A and 8C (adjusted p < 0.01, log2FC > 2). **e**, Genome coverage of 8A peaks that were not present in 4C or 8C embryos. **f**, Clustering (k=7) of TSS based on normalized H3K27ac signal (CPM) in 2C, 4C, 8C, and 8A embryos. **g**, Genome coverage of peaks gained or lost during the 4- to 8-cell transition, subcategorized based on whether they were present in 8A embryos. Peaks classified as either genic (overlapping gene bodies or 2Kb promoters) or intergenic.

**Extended Figure 8.**
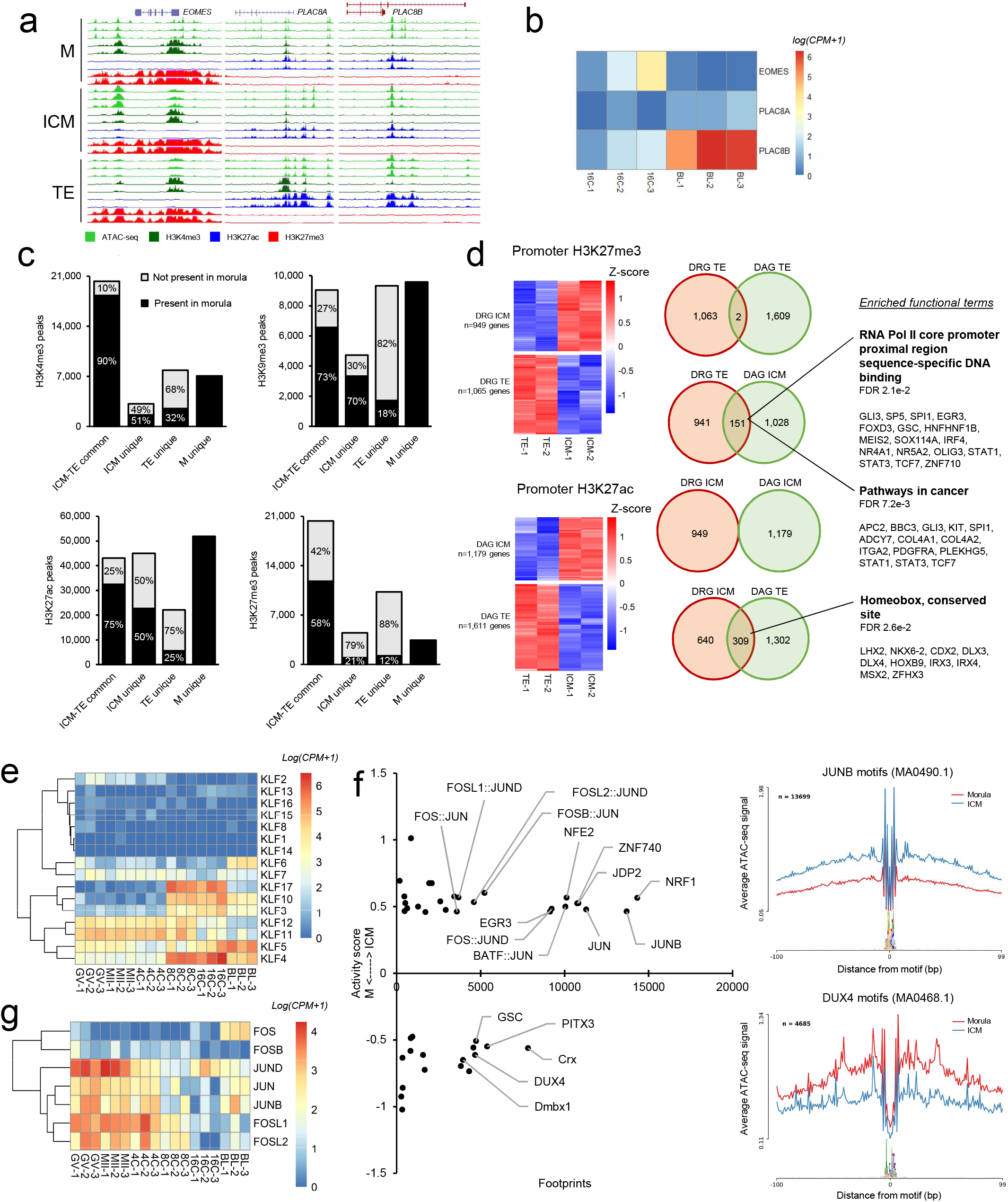
Epigenetic and transcription factor regulation of blastocyst lineages. **a**, Epigenetic profiles of *EOMES* (mouse-specific TE marker) and *PLAC8* (human-specific TE-marker). **b**, Normalized expression (CPM) of *EOMES* and *PLAC8*. **c**, Genome coverage of peaks unique to ICM, TE, or M, or shared in common between ICM and TE. Peaks categorized based on presence or absence in morula. **d**, Differentially regulated genes (adjusted p<0.05, log2FC>1) in ICM and TE based on H3K27me3 and H3K27me3 signal in 2Kb promoters. Enriched functional terms and corresponding genes reported for genes that were differentially activated (DAG) in one cell type and differentially repressed (DRG) in the other. **e**, Normalized expression of KLFs. **f**, TFs with differential activity (p<0.05) in ICM compared to M, based on footprinting analysis of ATAC-seq data. As examples, average signal shown at JUNB and DUX4 footprints. **g**, Normalized expression of JUN and FOS factors.

**Extended Figure 9.**
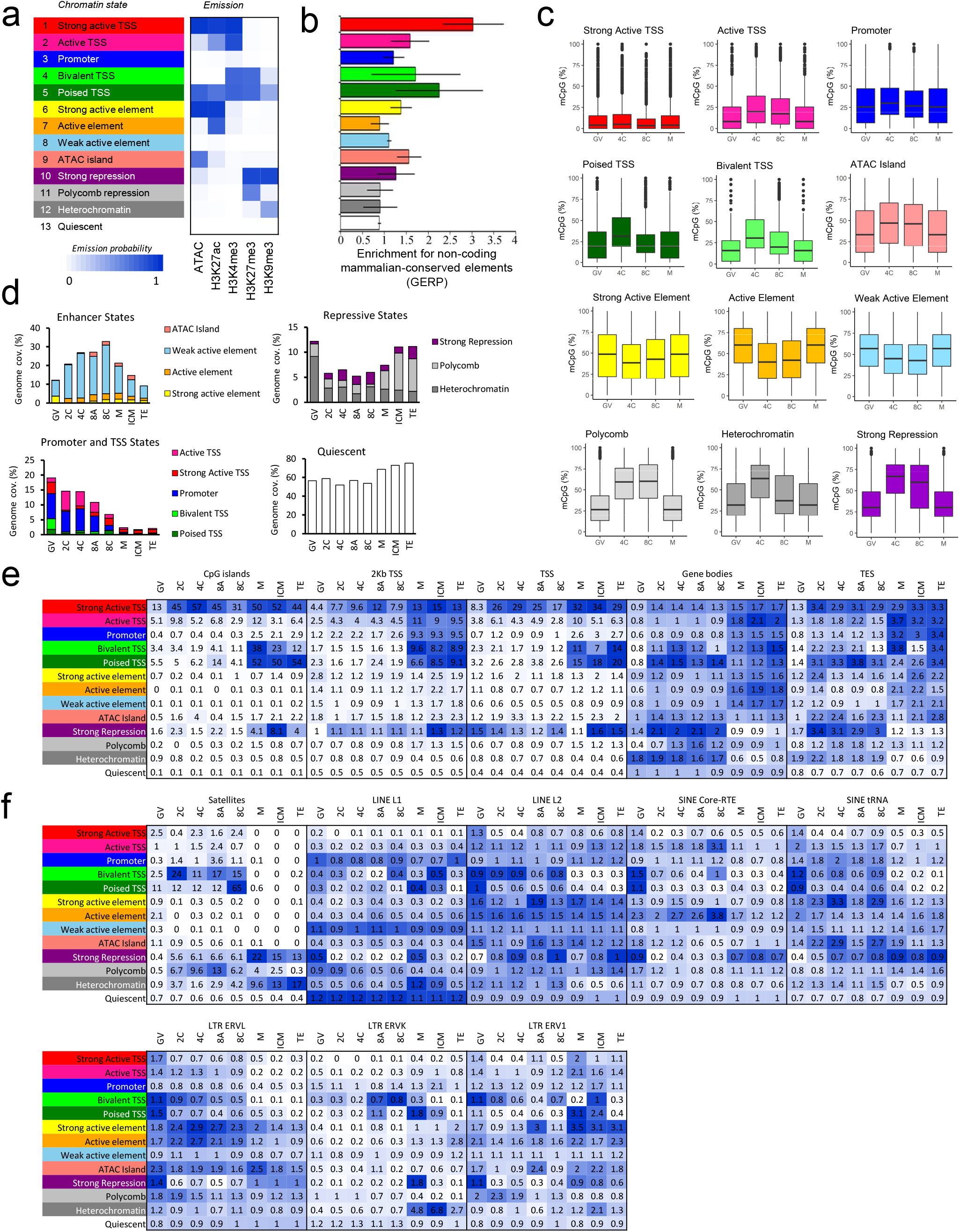
Characterization of the 13-state chromatin model. **a**, Chromatin state predictions based on accessibility and histone modification profiles. Emission probabilities indicate the likelihood of a given mark occurring in a given state. States were manually annotated based on **b**, sequence conservation, **c**, CpG methylation status, **d**, genome coverage, **e**, fold enrichment of overlap with CpG islands, 2Kb regions upstream of TSS, TSS, gene bodies, and TES, and **f**, fold enrichment of overlap with with satellite repeats, LINE, SINE, and LTR repetitive elements. For each genomic feature (e.g., “TSS”, “Satellites”, etc), fold enrichment values are shaded from white (lowest value) to blue (highest value).

**Extended Figure 10.**
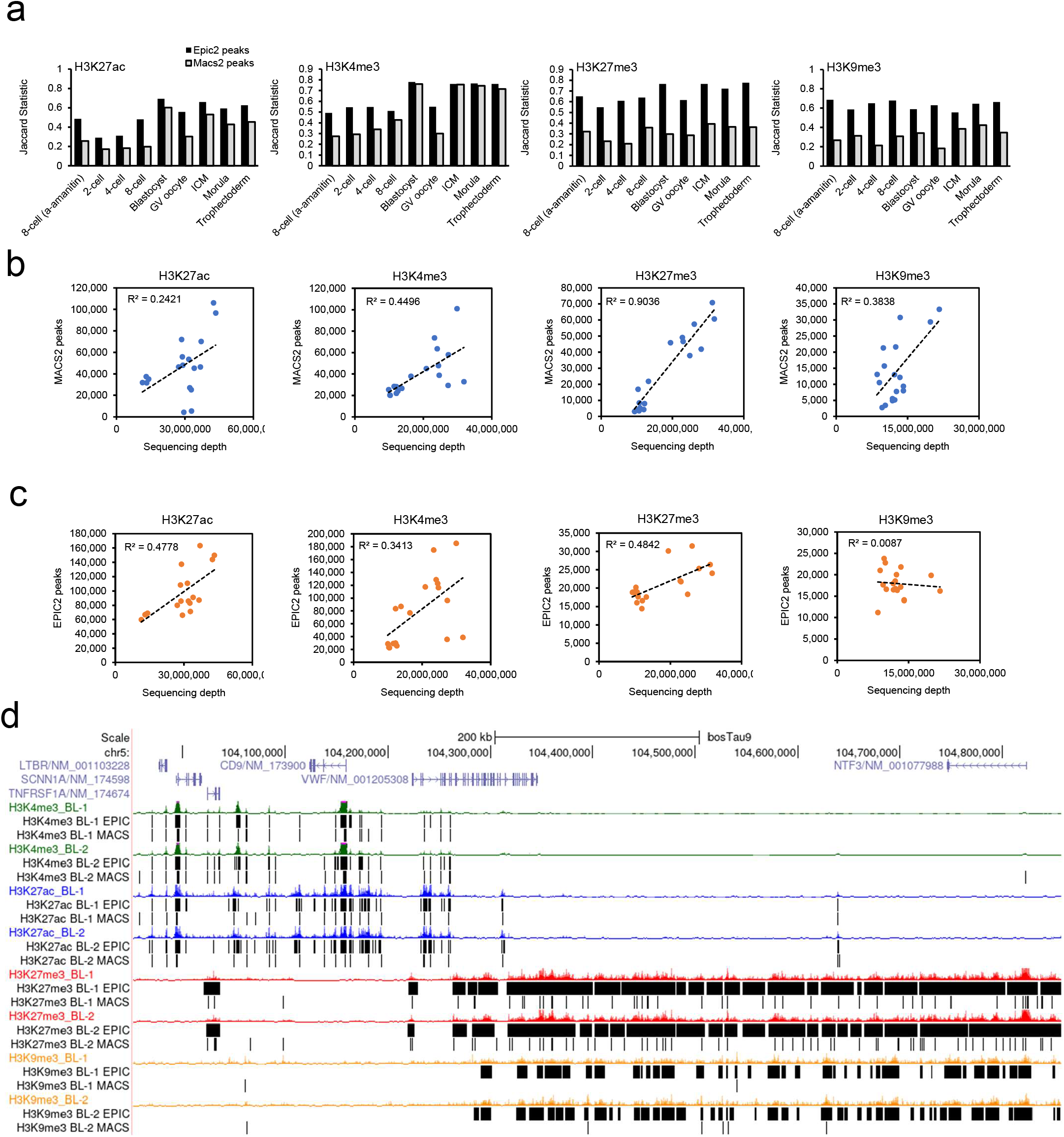
Comparison of peak callers Epic2 and Macs2. **a**, Overlap of peaks identified in biological replicates for each stage and mark, summarized using the Jaccard statistic, which measures the base-pair overlap between two peak sets, ranging from 0 (no overlap) to 1 (complete overlap). **b**, Correlation between sequencing depth (informative read alignments) and number of peaks called with either Epic2 or Macs2. **c**, Gene track view of Macs2 and Epic2 peaks called from CUT&Tag libraries generated from whole blastocysts for each mark.

**Extended Table 1.** Sequencing depth of H3K4me3 libraries. Aligned rate indicates percent of raw reads that were aligned, and duplicate rate indicates percent of aligned reads that were duplicates.

| Sample | Raw reads | Aligned reads | Aligned rate | Duplicate rate | Informative reads |
| --- | --- | --- | --- | --- | --- |
| K4me3_GV_1 | 50,069,438 | 48,936,121 | 97.86% | 31.54% | 23,279,878 |
| K4me3_GV_2 | 56,090,746 | 54,720,209 | 97.67% | 23.90% | 29,877,099 |
| K4me3_2C_1 | 54,682,218 | 53,306,796 | 97.62% | 25.30% | 24,432,941 |
| K4me3_2C_2 | 56,252,640 | 54,432,206 | 96.90% | 27.95% | 24,642,539 |
| K4me3_4C_1 | 51,121,482 | 49,844,161 | 97.62% | 25.95% | 20,853,662 |
| K4me3_4C_2 | 60,815,756 | 59,179,974 | 97.41% | 31.44% | 24,055,045 |
| K4me3_8C_1 | 101,667,142 | 96,309,699 | 94.86% | 58.93% | 16,378,114 |
| K4me3_8C_2 | 120,883,314 | 116,951,909 | 96.87% | 50.48% | 27,311,368 |
| K4me3_M_1 | 216,687,670 | 191,033,442 | 88.29% | 88.05% | 10,608,576 |
| K4me3_M_2 | 133,649,466 | 126,280,592 | 94.59% | 90.06% | 6,653,416 |
| K4me3_BL_1 | 102,268,744 | 99,167,118 | 97.09% | 69.30% | 27,227,953 |
| K4me3_BL_2 | 115,380,444 | 111,831,808 | 97.08% | 68.45% | 31,850,388 |
| K4me3_ICM_1 | 71,289,214 | 68,646,465 | 96.41% | 81.60% | 10,334,772 |
| K4me3_ICM_2 | 85,831,946 | 82,519,093 | 96.24% | 84.53% | 10,220,680 |
| K4me3_TE_1 | 166,083,630 | 86,267,784 | 96.93% | 72.04% | 39,586,455 |
| K4me3_TE_2 | 96,759,778 | 94,157,397 | 97.43% | 83.77% | 12,531,898 |
| K4me3_8A_1 | 44,611,340 | 43,313,882 | 97.24% | 33.81% | 13,755,559 |
| K4me3_8A_2 | 37,420,558 | 36,348,572 | 97.27% | 31.49% | 12,147,097 |

**Extended Table 2.** Sequencing depth of H3K27ac libraries. Aligned rate indicates percent of raw reads that were aligned, and duplicate rate indicates percent of aligned reads that were duplicates.

| Sample | Raw reads | Aligned reads | Aligned rate | Duplicate rate | Informative reads |
| --- | --- | --- | --- | --- | --- |
| K27ac_GV_1 | 64,275,194 | 62,432,336 | 97.26% | 28.51% | 37,065,865 |
| K27ac_GV_2 | 51,033,960 | 49,650,960 | 97.39% | 28.37% | 28,984,729 |
| K27ac_2C_1 | 46,507,932 | 45,451,802 | 97.85% | 17.29% | 29,427,433 |
| K27ac_2C_2 | 52,443,900 | 51,198,343 | 97.74% | 17.83% | 32,847,725 |
| K27ac_4C_1 | 51,608,892 | 50,420,909 | 97.81% | 12.52% | 34,023,225 |
| K27ac_4C_2 | 55,009,274 | 53,701,820 | 97.75% | 11.94% | 36,669,646 |
| K27ac_8C_1 | 52,145,308 | 51,072,871 | 98.08% | 21.76% | 32,682,964 |
| K27ac_8C_2 | 51,385,188 | 50,324,650 | 98.07% | 21.75% | 32,037,484 |
| K27ac_M_1 | 61,733,190 | 60,491,329 | 98.09% | 22.64% | 42,491,649 |
| K27ac_M_2 | 47,320,940 | 45,812,238 | 96.95% | 30.59% | 28,477,139 |
| K27ac_BL_1 | 38,323,414 | 37,524,483 | 98.03% | 63.76% | 11,352,801 |
| K27ac_BL_2 | 25,097,988 | 24,570,946 | 98.02% | 64.24% | 7,154,955 |
| K27ac_ICM_1 | 51,547,108 | 50,204,895 | 97.51% | 35.17% | 28,762,027 |
| K27ac_ICM_2 | 49,729,436 | 48,502,939 | 97.64% | 36.17% | 27,115,660 |
| K27ac_TE_1 | 62,425,412 | 59,680,774 | 95.72% | 72.39% | 13,194,360 |
| K27ac_TE_2 | 64,096,642 | 62,012,511 | 96.84% | 71.87% | 14,077,927 |
| K27ac_8A_1 | 78,096,352 | 76,112,532 | 97.61% | 20.99% | 43,426,169 |
| K27ac_8A_2 | 53,995,342 | 52,762,232 | 97.84% | 17.22% | 31,607,792 |

**Extended Table 3.** Sequencing depth of H3K27me3 libraries. Aligned rate indicates percent of raw reads that were aligned, and duplicate rate indicates percent of aligned reads that were duplicates.

| Sample | Raw reads | Aligned reads | Aligned rate | Duplicate rate | Informative reads |
| --- | --- | --- | --- | --- | --- |
| K27me3_GV_1 | 68,548,202 | 66,446,589 | 97.05% | 36.64% | 22,762,608 |
| K27me3_GV_2 | 63,585,900 | 61,436,688 | 96.74% | 38.30% | 19,411,346 |
| K27me3_2C_1 | 46,050,212 | 44,483,360 | 96.72% | 50.80% | 9,330,224 |
| K27me3_2C_2 | 61,349,468 | 59,280,917 | 96.75% | 63.89% | 7,014,178 |
| K27me3_4C_1 | 51,521,262 | 49,970,663 | 97.10% | 47.36% | 12,144,222 |
| K27me3_4C_2 | 83,128,130 | 80,518,620 | 96.95% | 61.41% | 10,904,547 |
| K27me3_8C_1 | 93,003,634 | 89,585,287 | 96.41% | 57.90% | 10,608,241 |
| K27me3_8C_2 | 139,880,648 | 134,822,780 | 96.49% | 63.87% | 11,947,092 |
| K27me3_M_1 | 94,392,770 | 91,056,979 | 96.60% | 56.08% | 13,238,181 |
| K27me3_M_2 | 52,867,942 | 51,479,776 | 97.50% | 44.98% | 10,309,192 |
| K27me3_BL_1 | 44,436,330 | 43,194,312 | 97.30% | 21.10% | 22,944,311 |
| K27me3_BL_2 | 60,798,324 | 58,979,181 | 97.14% | 23.20% | 31,719,305 |
| K27me3_ICM_1 | 56,806,354 | 55,137,389 | 97.19% | 25.14% | 28,028,647 |
| K27me3_ICM_2 | 48,598,228 | 47,348,993 | 97.53% | 24.18% | 24,800,117 |
| K27me3_TE_1 | 56,300,916 | 54,723,374 | 97.32% | 29.25% | 26,079,844 |
| K27me3_TE_2 | 70,886,866 | 69,020,203 | 97.49% | 33.24% | 31,123,330 |
| K27me3_8A_1 | 179,025,096 | 169,241,102 | 94.67% | 74.21% | 9,741,798 |
| K27me3_8A_2 | 137,951,488 | 131,145,350 | 95.20% | 70.10% | 10,257,664 |

**Extended Table 4.**
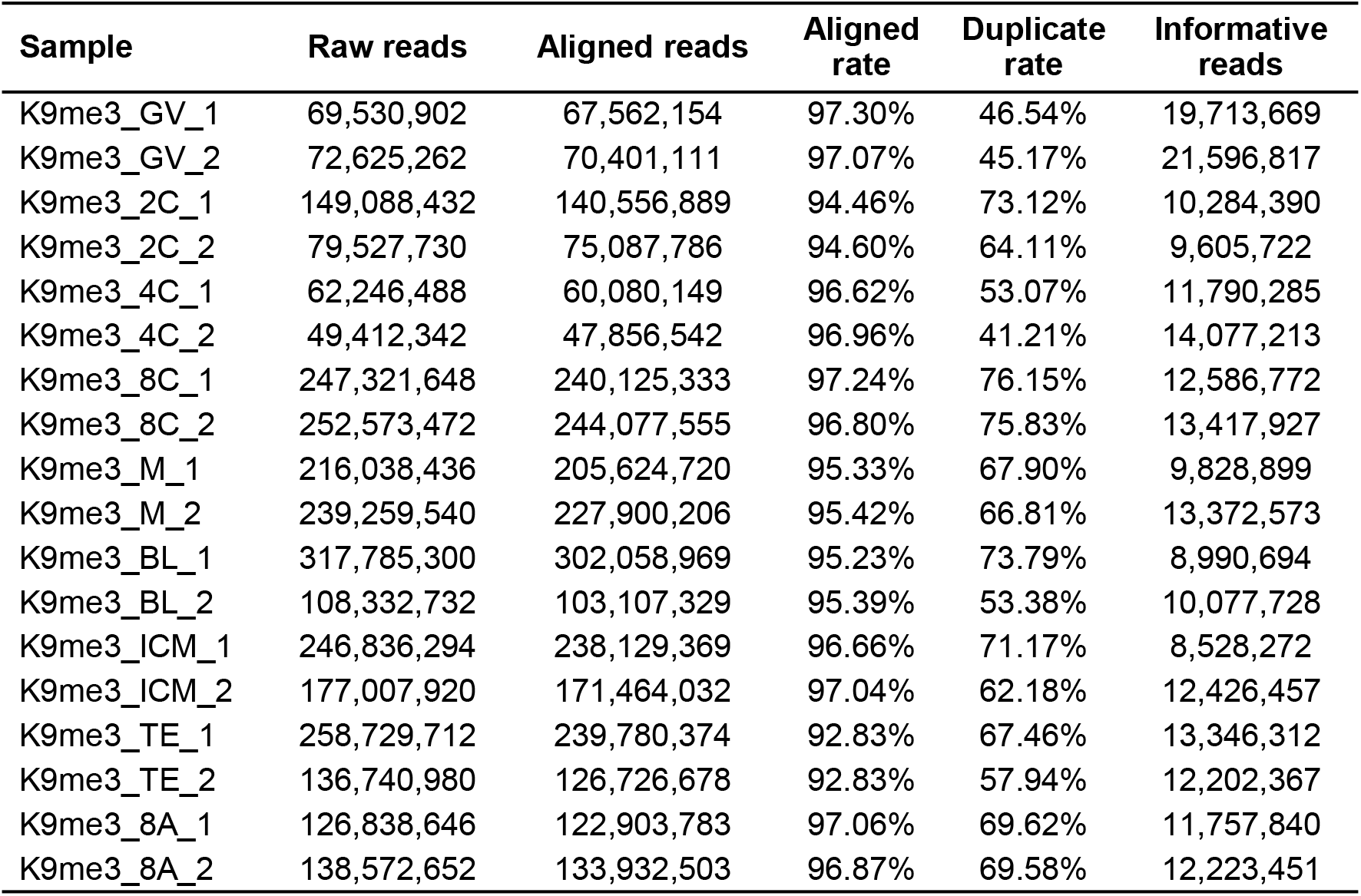
Sequencing depth of H3K9me3 libraries. Aligned rate indicates percent of raw reads that were aligned, and duplicate rate indicates percent of aligned reads that were duplicates. PMDs.

**Extended Table 5.** Sequencing depth of ATAC-seq libraries. Aligned rate indicates percent of raw reads that were aligned, and duplicate rate indicates percent of aligned reads that were duplicates.

| Sample | Raw reads | Aligned reads | Aligned rate | Duplicate rate | Informative reads |
| --- | --- | --- | --- | --- | --- |
| ATAC_BL_1 | 61,226,258 | 51,985,747 | 85.03% | 12.25% | 38,791,641 |
| ATAC_BL_2 | 73,624,292 | 68,749,779 | 93.52% | 10.45% | 52,313,227 |
| ATAC_BL_3 | 111,446,012 | 96,981,910 | 87.17% | 11.32% | 73,433,906 |
| ATAC_ICM_1 | 59,628,982 | 50,871,935 | 85.47% | 10.72% | 39,663,063 |
| ATAC_ICM_2 | 47,690,862 | 42,883,828 | 90.03% | 8.24% | 33,967,255 |
| ATAC_ICM_3 | 51,839,666 | 46,143,818 | 89.11% | 9.33% | 36,092,796 |
| ATAC_TE_1 | 53,273,238 | 39,682,695 | 74.56% | 20.57% | 26,968,756 |
| ATAC_TE_2 | 65,835,150 | 49,114,692 | 74.69% | 21.09% | 33,182,182 |
| ATAC_TE_3 | 77,640,516 | 57,689,575 | 74.40% | 24.22% | 37,439,938 |

**Extended Table 6.**
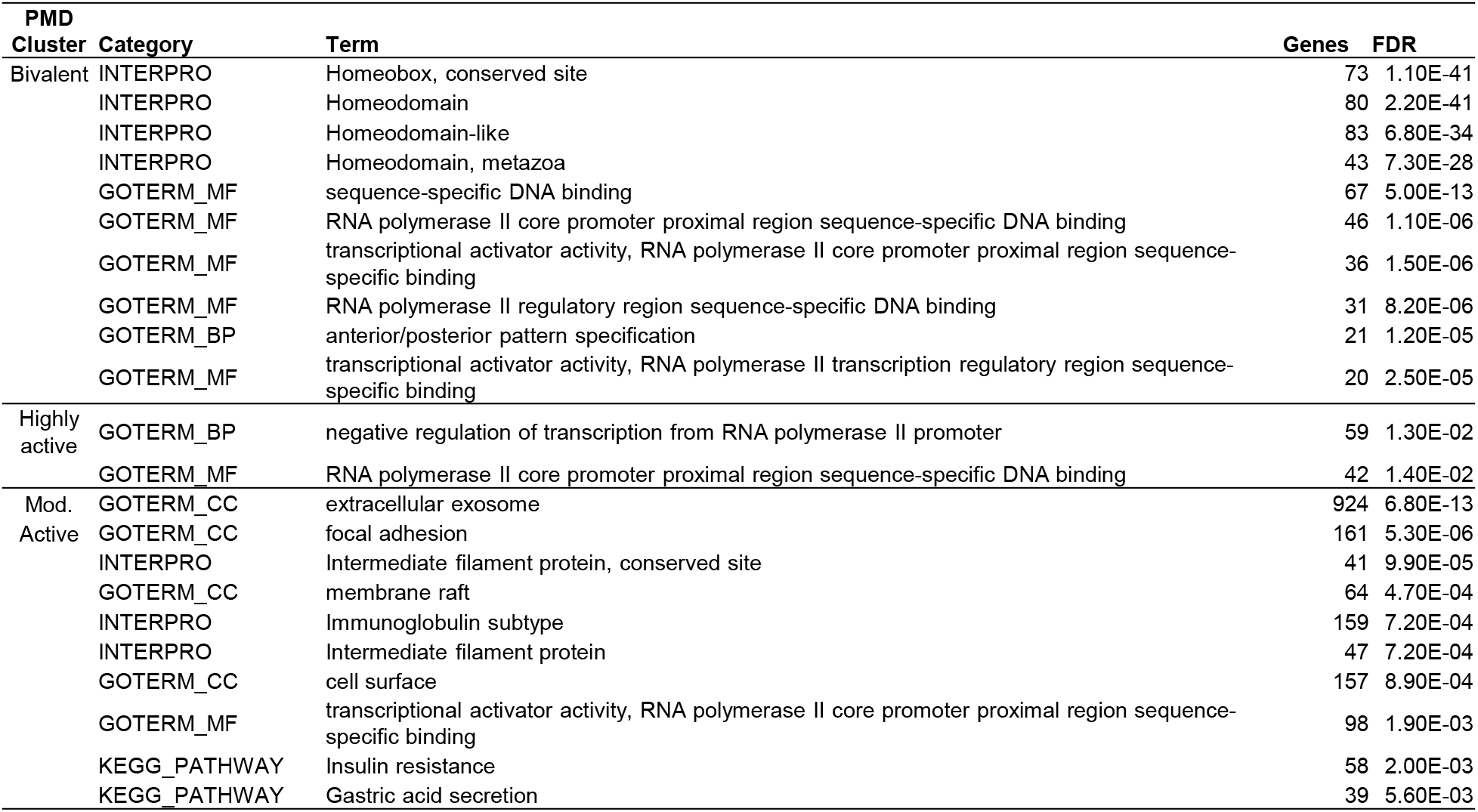
Functional annotation of genes marked by PMD clusters. Top 10 terms for each cluster, based on FDR, are reported when available. No gene ontology (GO), Interpro, or KEGG pathways were enriched in genes marked by quiescent PMDs.

**Extended Table 7.** For each epigenetic mark, number of peaks that were gained, lost, or retained during the 4- to 8-cell transition. Gained peaks were found in 8-cell (8C) embryos, but not 4-cell (4C). Lost peaks were found in 4C, but not 8C. Retained peaks were found in both 4C and 8C embryos. Peaks are further sub-categorized based on whether they were present or absent in 8-cell embryos cultured in the presence of a-amanitin (8A).

|  | ATAC-seq |  | H3K4me3 |  | H3K27ac |  | H3K27me3 |  | H3K9me3 |  |
| --- | --- | --- | --- | --- | --- | --- | --- | --- | --- | --- |
| **Gained peaks (4C to 8C)** | 68,035 |  | 2,943 |  | 18,969 |  | 3,588 |  | 3,087 |  |
| Present in 8A | 13,631 | 20% | 809 | 27% | 4,356 | 23% | 1,711 | 48% | 1,137 | 37% |
| Not present in 8A | 54,404 | 80% | 2,134 | 73% | 14,613 | 77% | 1,877 | 52% | 1,950 | 63% |
| **Lost peaks (4C to 8C)** | 1,892 |  | 7,924 |  | 10,889 |  | 4,237 |  | 2,386 |  |
| Not present in 8A | 435 | 23% | 5,056 | 64% | 2,913 | 27% | 2,100 | 50% | 969 | 41% |
| Present in 8A | 1,457 | 77% | 2,868 | 36% | 7,976 | 73% | 2,137 | 50% | 1,417 | 59% |
| **Retained peaks (4C to 8C)** | 6,278 |  | 23,717 |  | 12,060 |  | 10,639 |  | 11,572 |  |
| Present in 8A | 5,931 | 94% | 21,876 | 92% | 10,513 | 87% | 9,670 | 91% | 10,596 | 92% |
| Not present in 8A | 347 | 6% | 1,841 | 8% | 1,547 | 13% | 969 | 9% | 976 | 8% |

**Extended Table 8.** Functional annotation of genes belonging to transcript clusters identified based on H3K27ac signal at TSS in 2-cell, 4-cell, 8-cell, and 8-cell embryos treated with a-amanitin (Extended Fig. 17b). Top 10 terms, based on FDR, are reported for clusters 1-5.

| Cluster | Category | Term | Genes | % | FDR |
| --- | --- | --- | --- | --- | --- |
| 1 | GOTERM_CC | nucleus | 274 | 32 | 3.10E-27 |
|  | GOTERM_MF | RNA polymerase II transcription factor activity, sequence-specific DNA binding | 106 | 12 | 1.80E-20 |
|  | GOTERM_MF | RNA polymerase II core promoter proximal region sequence-specific DNA binding | 109 | 13 | 6.80E-20 |
|  | GOTERM_BP | regulation of transcription from RNA polymerase II promoter | 132 | 15 | 1.90E-16 |
|  | INTERPRO | Homeodomain-like | 48 | 6 | 7.70E-16 |
|  | INTERPRO | Homeobox, conserved site | 37 | 4 | 2.90E-15 |
|  | INTERPRO | Homeodomain | 40 | 5 | 3.00E-14 |
|  | GOTERM_CC | nucleoplasm | 142 | 17 | 1.50E-10 |
|  | GOTERM_MF | transcriptional activator activity, RNA polymerase II transcription regulatory region sequence-specific binding | 47 | 6 | 3.50E-09 |
|  | GOTERM_MF | mRNA binding | 29 | 3 | 2.80E-07 |
| 2 | GOTERM_CC | cytosol | 620 | 21 | 9.70E-46 |
|  | GOTERM_CC | nucleus | 768 | 26 | 2.80E-38 |
|  | GOTERM_CC | nucleoplasm | 477 | 16 | 4.40E-37 |
|  | GOTERM_CC | nucleolus | 180 | 6 | 1.90E-23 |
|  | GOTERM_MF | RNA binding | 186 | 6 | 2.20E-22 |
|  | GOTERM_CC | cytoplasm | 653 | 22 | 9.80E-18 |
|  | GOTERM_CC | mitochondrion | 212 | 7 | 4.40E-16 |
|  | GOTERM_CC | centrosome | 117 | 4 | 3.20E-14 |
|  | GOTERM_CC | nuclear speck | 94 | 3 | 2.70E-11 |
|  | GOTERM_MF | ATP binding | 262 | 9 | 6.10E-10 |
| 3 | GOTERM_MF | RNA polymerase II transcription factor activity, sequence-specific DNA binding | 204 | 8 | 2.40E-15 |
|  | GOTERM_MF | RNA polymerase II core promoter proximal region sequence-specific DNA binding | 212 | 8 | 1.40E-14 |
|  | INTERPRO | Homeodomain | 70 | 3 | 2.20E-12 |
|  | INTERPRO | Homeobox, conserved site | 59 | 2 | 6.50E-12 |
|  | INTERPRO | Homeodomain-like | 77 | 3 | 2.00E-10 |
|  | GOTERM_BP | regulation of transcription from RNA polymerase II promoter | 268 | 10 | 2.80E-09 |
|  | GOTERM_CC | nucleus | 571 | 21 | 2.70E-08 |
|  | GOTERM_CC | nucleoplasm | 327 | 12 | 1.00E-06 |
|  | GOTERM_CC | cytoplasm | 533 | 20 | 1.10E-06 |
| 4 | GOTERM_CC | nucleus | 535 | 23 | 1.60E-20 |
|  | GOTERM_MF | RNA polymerase II core promoter proximal region sequence-specific DNA binding | 181 | 8 | 1.20E-14 |
|  | GOTERM_MF | RNA polymerase II transcription factor activity, sequence-specific DNA binding | 167 | 7 | 4.70E-13 |
|  | GOTERM_CC | nucleoplasm | 302 | 13 | 1.50E-12 |
|  | GOTERM_CC | cytoplasm | 477 | 21 | 2.40E-12 |
|  | GOTERM_CC | cytosol | 364 | 16 | 2.00E-09 |
|  | GOTERM_MF | transcriptional activator activity, RNA polymerase II transcription regulatory region sequence-specific binding | 80 | 3 | 1.60E-08 |
|  | GOTERM_BP | regulation of transcription from RNA polymerase II promoter | 219 | 9 | 1.20E-07 |
|  | GOTERM_MF | protein kinase binding | 65 | 3 | 1.80E-05 |
|  | GOTERM_MF | transcriptional repressor activity, RNA polymerase II transcription regulatory region sequence-specific binding | 54 | 2 | 2.20E-05 |
| 5 | GOTERM_CC | cytoplasm | 1097 | 21 | 7.70E-25 |
|  | GOTERM_CC | cytosol | 854 | 16 | 5.30E-23 |
|  | GOTERM_MF | ATP binding | 410 | 8 | 2.50E-08 |
|  | GOTERM_CC | nucleoplasm | 593 | 11 | 5.40E-08 |
|  | GOTERM_CC | Golgi apparatus | 231 | 4 | 4.20E-06 |
|  | GOTERM_CC | mitochondrion | 279 | 5 | 4.20E-06 |
|  | KEGG_PATHWAY | Metabolic pathways | 455 | 9 | 1.30E-05 |
|  | GOTERM_CC | membrane | 249 | 5 | 3.00E-05 |
|  | GOTERM_MF | protein homodimerization activity | 173 | 3 | 5.30E-05 |
|  | GOTERM_CC | nucleolus | 197 | 4 | 7.70E-05 |

**Extended Table 9.** Functional enrichment of genes marked by each peak set. Top 10 terms, including gene ontology terms (GOTERM) for cellular compartment (CC), molecular function (MF), and INTERPRO terms, based on FDR, are reported.

| Genic H3K27ac peaks gained from 4C to 8C |  |  | Peak not detected in 8A<br>(4,424 genes) |  |
| --- | --- | --- | --- | --- |
|  | Term |  | FDR | Genes |
| 1 | GOTERM_CC | cytosol | 8.70E-23 | 709 |
| 2 | GOTERM_MF | ATP binding | 1.30E-17 | 383 |
| 3 | GOTERM_CC | cytoplasm | 5.20E-17 | 864 |
| 4 | GOTERM_CC | nucleoplasm | 6.20E-16 | 528 |
| 5 | GOTERM_CC | nucleus | 3.20E-12 | 873 |
| 6 | GOTERM_MF | metal ion binding | 1.30E-11 | 396 |
| 7 | INTERPRO | Pleckstrin homology-like domain | 1.00E-09 | 133 |
| 8 | INTERPRO | P-loop containing nucleoside triphosphate hydrolase | 2.70E-08 | 241 |
| 9 | GOTERM_MF | actin binding | 2.30E-07 | 80 |
| 10 | INTERPRO | Armadillo-type fold | 1.00E-06 | 109 |

| Genic H3K27ac peaks lost from 4C to 8C |  |  | Peak detected in 8A<br>(2,616 genes) |  |
| --- | --- | --- | --- | --- |
|  | Term |  | FDR | Genes |
| 1 | GOTERM_MF | calcium ion binding | 3.70E-09 | 119 |
| 2 | KEGG_PATHWAY | Pathways in cancer | 9.30E-06 | 95 |
| 3 | KEGG_PATHWAY | MAPK signaling pathway | 9.30E-06 | 60 |
| 4 | KEGG_PATHWAY | Calcium signaling pathway | 9.30E-06 | 54 |
| 5 | KEGG_PATHWAY | Oxytocin signaling pathway | 1.70E-05 | 38 |
| 6 | KEGG_PATHWAY | Ras signaling pathway | 2.60E-05 | 51 |
| 7 | GOTERM_CC | extracellular matrix | 4.20E-05 | 46 |
| 8 | GOTERM_CC | voltage-gated potassium channel complex | 4.20E-05 | 22 |
| 9 | GOTERM_CC | integral component of plasma membrane | 6.90E-05 | 131 |
| 10 | GOTERM_CC | apical plasma membrane | 6.90E-05 | 48 |

| Genic H3K27ac peaks gained from 4C to 8C |  |  | Peak detected in 8A<br>(2,974 genes) |  |
| --- | --- | --- | --- | --- |
|  | Term |  | FDR | Genes |
| 1 | GOTERM_CC | nucleus | 1.30E-31 | 719 |
| 2 | GOTERM_CC | nucleoplasm | 1.20E-28 | 437 |
| 3 | GOTERM_CC | cytoplasm | 9.10E-28 | 675 |
| 4 | GOTERM_CC | cytosol | 4.00E-23 | 523 |
| 5 | GOTERM_CC | nucleolus | 3.60E-13 | 150 |
| 6 | GOTERM_MF | ATP binding | 3.60E-11 | 265 |
| 7 | GOTERM_CC | nuclear speck | 1.40E-10 | 90 |
| 8 | INTERPRO | Armadillo-type fold | 1.60E-08 | 89 |
| 9 | GOTERM_CC | centrosome | 9.20E-08 | 97 |
| 10 | INTERPRO | P-loop containing nucleoside triphosphate hydrolase | 1.20E-07 | 175 |

| Genic H3K27ac peaks lost from 4C to 8C |  |  | Peak not detected in 8A<br>(827 genes) |  |
| --- | --- | --- | --- | --- |
|  | Term |  | FDR | Genes |
| 1 | KEGG_PATHWAY | Calcium signaling pathway | 3.60E-04 | 25 |
| 2 | GOTERM_CC | integral component of plasma membrane | 1.30E-03 | 55 |
| 3 | GOTERM_CC | postsynaptic membrane | 1.50E-03 | 15 |
| 4 | KEGG_PATHWAY | Cholinergic synapse | 1.80E-03 | 15 |
| 5 | KEGG_PATHWAY | Adrenergic signaling in cardiomyocytes | 1.80E-03 | 17 |
| 6 | KEGG_PATHWAY | Aldosterone synthesis and secretion | 3.30E-03 | 13 |
| 7 | KEGG_PATHWAY | Neuroactive ligand-receptor interaction | 3.60E-03 | 28 |
| 8 | KEGG_PATHWAY | cGMP-PKG signaling pathway | 4.60E-03 | 17 |
| 9 | KEGG_PATHWAY | Vascular smooth muscle contraction | 5.30E-03 | 15 |
| 10 | GOTERM_CC | membrane raft | 8.10E-03 | 15 |

**Extended Table 10.** Enriched gene ontology terms for differentially repressed (higher H3K27me3) and differentially activated (higher H3K27ac) gene sets in ICM and TE. Significant gene ontology (GOTERM) terms (FDR<0.05, MF; molecular function, BP; biological process) are reported.

| Category | Term | Genes | FDR |
| --- | --- | --- | --- |
| GOTERM_MF | RNA polymerase II transcription factor activity, sequence-specific DNA binding | 98 | 1.80E-19 |
| GOTERM_MF | RNA polymerase II core promoter proximal region sequence-specific DNA binding | 98 | 1.60E-17 |
| GOTERM_MF | transcriptional activator activity, RNA polymerase II transcription regulatory region sequence-specific binding | 50 | 1.40E-12 |
| GOTERM_BP | regulation of transcription from RNA polymerase II promoter | 110 | 1.90E-12 |
| GOTERM_BP | embryonic skeletal system morphogenesis | 14 | 2.60E-07 |
| GOTERM_MF | transcriptional repressor activity, RNA polymerase II transcription regulatory region sequence-specific binding | 28 | 1.00E-04 |
| GOTERM_MF | RNA polymerase II regulatory region sequence-specific DNA binding | 21 | 9.60E-04 |
| GOTERM_BP | epithelial tube branching involved in lung morphogenesis | 7 | 2.00E-03 |
| GOTERM_BP | anterior/posterior pattern specification | 15 | 2.00E-03 |
| GOTERM_MF | glycine N-acyltransferase activity | 5 | 5.00E-03 |
| GOTERM_BP | inner ear morphogenesis | 10 | 5.10E-03 |
| GOTERM_BP | proximal/distal pattern formation | 8 | 5.10E-03 |
| GOTERM_BP | negative regulation of transcription from RNA polymerase II promoter | 41 | 6.60E-03 |
| GOTERM_MF | steroid hydroxylase activity | 8 | 1.60E-02 |
| GOTERM_MF | G-protein coupled peptide receptor activity | 8 | 1.60E-02 |
| GOTERM_BP | positive regulation of transcription from RNA polymerase II promoter | 47 | 1.70E-02 |
| GOTERM_BP | regulation of neurogenesis | 8 | 1.80E-02 |
| GOTERM_BP | cell differentiation | 30 | 3.80E-02 |

*Genes differentially marked by H3K27ac in ICM*
| Category | Term | Genes | FDR |
| --- | --- | --- | --- |
| GOTERM_MF | DNA binding | 63 | 6.40E-03 |
| GOTERM_MF | transcription factor activity, sequence-specific DNA binding | 28 | 6.40E-03 |
| GOTERM_MF | RNA polymerase II core promoter proximal region sequence-specific DNA binding | 77 | 6.40E-03 |
| GOTERM_MF | ATP binding | 93 | 4.30E-02 |

*Genes differentially marked by H3K27me3 in TE*
| Category | Term | Genes | FDR |
| --- | --- | --- | --- |
| GOTERM_MF | calcium ion binding | 52 | 6.00E-04 |
| GOTERM_BP | defense response | 10 | 7.10E-03 |

*Genes differentially marked by H3K27ac in TE*
| Category | Term | Genes | FDR |
| --- | --- | --- | --- |
| GOTERM_BP | cholesterol biosynthetic process | 11 | 1.50E-02 |
| GOTERM_MF | phosphatidylinositol binding | 18 | 2.30E-02 |

Differentially marked by H3K27me3 in ICM

**Extended Table 11.**
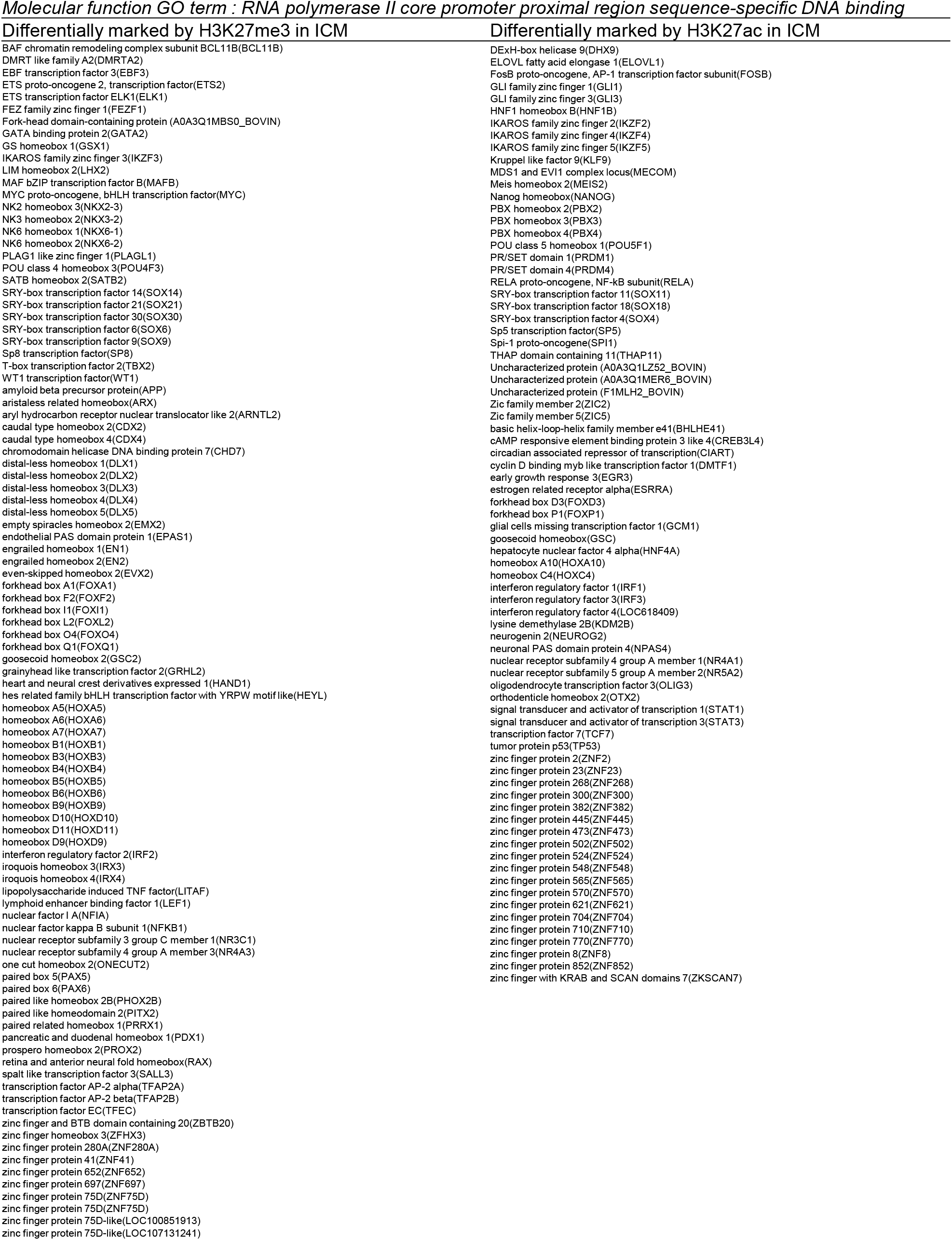
Differentially repressed and differentially activated genes in ICM that correspond to the enriched gene ontology term “RNA polymerase II core promoter proximal region sequence-specific DNA binding.”

**Extended Table 12.** Parameters used for peak calling with EPIC2 and MACS2.

| *Epigenetic mark* | *EPIC2* | *MACS2* |
| --- | --- | --- |
| H3K4me3 (narrow) | --effective-genome-fraction 0.7<br>-bin 50<br>-g 1<br>-e 1 | -q 0.05<br>-B |
| H3K27ac (narrow) | --effective-genome-fraction 0.7<br>-bin 50<br>-g 1<br>-e 1 | -q 0.05<br>-B |
| H3K4me3 (broad) | --effective-genome-fraction 0.7<br>-bin 200<br>-g 5<br>-e 1 |  |
| H3K27ac (broad) | --effective-genome-fraction 0.7<br>-bin 200<br>-g 5<br>-e 1 | - |
| H3K27me3 | --effective-genome-fraction 0.7<br>-bin 200<br>-g 5<br>-e 1 | -q 0.05<br>-B<br>--broad |
| H3K9me3 | --effective-genome-fraction 0.7<br>-bin 200<br>-g 5<br>-e 1 | -q 0.05<br>-B<br>--broad |
| ATAC-seq | --effective-genome-fraction 0.7<br>-bin 50<br>-g 1<br>-e 1 | -q 0.05<br>--shift -100<br>--extsize -200<br>-B |

## Notes

### Competing Interest Statement

The authors have declared no competing interest.

## References

1. Schultz, R. M., Stein, P. & Svoboda, P. The oocyte-to-embryo transition in mouse: past, present, and future. Biol. Reprod. 99, 160–174 (2018).

2. Frum, T. & Ralston, A. Cell signaling and transcription factors regulating cell fate during formation of the mouse blastocyst. Trends Genet. 31, 402–410 (2015).

3. Schulz, K. N. & Harrison, M. M. Mechanisms regulating zygotic genome activation. Nat. Rev. Genet. 20, 221–234 (2019).

4. Jambhekar, A., Dhall, A. & Shi, Y. Roles and regulation of histone methylation in animal development. Nat. Rev. Mol. cell Biol. 20, 625–641 (2019).

5. Eckersley-Maslin, M. A., Alda-Catalinas, C. & Reik, W. Dynamics of the epigenetic landscape during the maternal-to-zygotic transition. Nat. Rev. Mol. Cell Biol. 19, 436–450 (2018).

6. Li, L. et al. Single-cell multi-omics sequencing of human early embryos. Nat. Cell Biol. 20, 847–858 (2018).

7. Wu, J. et al. Chromatin analysis in human early development reveals epigenetic transition during ZGA. (2018) doi:10.1038/s41586-018-0080-8.

8. Gao, L. et al. Chromatin Accessibility Landscape in Human Early Embryos and Its Association with Evolution. Cell 173, 248-259.e15 (2018).

9. Liu, L. et al. An integrated chromatin accessibility and transcriptome landscape of human pre-implantation embryos. Nat. Commun. 10, 364 (2019).

10. Lu, F. et al. Establishing Chromatin Regulatory Landscape during Mouse Preimplantation Development. Cell 165, 1375–1388 (2016).

11. Wu, J. et al. The landscape of accessible chromatin in mammalian preimplantation embryos. Nature 534, 652–657 (2016).

12. Halstead, M. M., Ma, X., Zhou, C., Schultz, R. M. & Ross, P. J. Chromatin remodeling in bovine embryos indicates species-specific regulation of genome activation. Nat. Commun. 11, 1–16 (2020).

13. Ming, H. et al. The landscape of accessible chromatin in bovine oocytes and early embryos. Epigenetics 16, 300–312 (2021).

14. Dahl, J. A. et al. Broad histone H3K4me3 domains in mouse oocytes modulate maternal-to-zygotic transition. Nature 537, 548–552 (2016).

15. Liu, X. et al. Distinct features of H3K4me3 and H3K27me3 chromatin domains in pre-implantation embryos. Nature 537, 558–562 (2016).

16. Zhang, B. et al. Allelic reprogramming of the histone modification H3K4me3 in early mammalian development. Nature 537, 553–557 (2016).

17. Zheng, H. et al. Resetting Epigenetic Memory by Reprogramming of Histone Modifications in Mammals. Mol. Cell 63, 1066–1079 (2016).

18. Wang, C. et al. Reprogramming of H3K9me3-dependent heterochromatin during mammalian embryo development. Nat. Cell Biol. 20, 620–631 (2018).

19. Xia, W. et al. Resetting histone modifications during human parental-to-zygotic transition. Science (80-.). 365, 353–360 (2019).

20. Lu, X. et al. Evolutionary epigenomic analyses in mammalian early embryos reveal species-specific innovations and conserved principles of imprinting. Sci. Adv. 7, eabi6178 (2021).

21. Zhang, W. et al. Maternal-biased H3K27me3 correlates with paternal-specific gene expression in the human morula. Genes Dev. 33, 382–387 (2019).

22. Kaya-Okur, H. S. et al. CUT&Tag for efficient epigenomic profiling of small samples and single cells. Nat. Commun. 10, 1–10 (2019).

23. Buenrostro, J. D., Giresi, P. G., Zaba, L. C., Chang, H. Y. & Greenleaf, W. J. Transposition of native chromatin for fast and sensitive epigenomic profiling of open chromatin, DNA-binding proteins and nucleosome position. Nat Methods 10, 1213–1218 (2013).

24. Canovas, S., Cibelli, J. B. & Ross, P. J. Jumonji domain-containing protein 3 regulates histone 3 lysine 27 methylation during bovine preimplantation development. Proc. Natl. Acad. Sci. U. S. A. 109, 2400–2405 (2012).

25. Ross, P. J. et al. Polycomb gene expression and histone H3 lysine 27 trimethylation changes during bovine preimplantation development. Reproduction 136, 777–785 (2008).

26. Zhou, C. et al. H3K27me3 is an epigenetic barrier while KDM6A overexpression improves nuclear reprogramming efficiency. FASEB J. 33, 4638–4652 (2019).

27. Skene, P. J. & Henikoff, S. CUT&RUN: Targeted in situ genome-wide profiling with high efficiency for low cell numbers. biorxiv 193219 (2017).

28. Ooga, M., Fulka, H., Hashimoto, S., Suzuki, M. G. & Aoki, F. Analysis of chromatin structure in mouse preimplantation embryos by fluorescent recovery after photobleaching. Epigenetics 11, 85–94 (2016).

29. Ahmed, K. et al. Global chromatin architecture reflects pluripotency and lineage commitment in the early mouse embryo. PLoS One 5, e10531–e10531 (2010).

30. Du, Z. et al. Allelic reprogramming of 3D chromatin architecture during early mammalian development. Nature 547, 232–235 (2017).

31. Wiekowski, M., Miranda, M. & DePamphilis, M. L. Requirements for promoter activity in mouse oocytes and embryos distinguish paternal pronuclei from maternal and zygotic nuclei. Dev. Biol. 159, 366–378 (1993).

32. Ivanova, E. et al. DNA methylation changes during preimplantation development reveal inter-species differences and reprogramming events at imprinted genes. Clin. Epigenetics 12, 1–18 (2020).

33. Greenberg, M. V. C. & Bourc’his, D. The diverse roles of DNA methylation in mammalian development and disease. Nat. Rev. Mol. cell Biol. 20, 590–607 (2019).

34. Rose, N. R. & Klose, R. J. Understanding the relationship between DNA methylation and histone lysine methylation. Biochim. Biophys. Acta (BBA)-Gene Regul. Mech. 1839, 1362–1372 (2014).

35. Bernstein, B. E. et al. A bivalent chromatin structure marks key developmental genes in embryonic stem cells. Cell 125, 315–326 (2006).

36. Xu, Q. et al. SETD2 regulates the maternal epigenome, genomic imprinting and embryonic development. Nat. Genet. 51, 844–856 (2019).

37. Bilodeau, S., Kagey, M. H., Frampton, G. M., Rahl, P. B. & Young, R. A. SetDB1 contributes to repression of genes encoding developmental regulators and maintenance of ES cell state. Genes Dev. 23, 2484–2489 (2009).

38. Alder, O. et al. Ring1B and Suv39h1 delineate distinct chromatin states at bivalent genes during early mouse lineage commitment. Development 137, 2483–2492 (2010).

39. Matsumura, Y. et al. H3K4/H3K9me3 bivalent chromatin domains targeted by lineage-specific DNA methylation pauses adipocyte differentiation. Mol. Cell 60, 584–596 (2015).

40. Hagarman, J. A., Motley, M. P., Kristjansdottir, K. & Soloway, P. D. Coordinate regulation of DNA methylation and H3K27me3 in mouse embryonic stem cells. PLoS One 8, e53880 (2013).

41. Chen, Z. & Zhang, Y. Maternal H3K27me3-dependent autosomal and X chromosome imprinting. Nat. Rev. Genet. 21, 555–571 (2020).

42. Local, A. et al. Identification of H3K4me1-associated proteins at mammalian enhancers. Nat. Genet. 50, 73–82 (2018).

43. Creyghton, M. P. et al. Histone H3K27ac separates active from poised enhancers and predicts developmental state. Proc Natl Acad Sci U S A 107, 21931–21936 (2010).

44. Rada-Iglesias, A. et al. A unique chromatin signature uncovers early developmental enhancers in humans. Nature 470, 279–283 (2011).

45. Zentner, G. E., Tesar, P. J. & Scacheri, P. C. Epigenetic signatures distinguish multiple classes of enhancers with distinct cellular functions. Genome Res. 21, 1273–1283 (2011).

46. Aoshima, K., Inoue, E., Sawa, H. & Okada, Y. Paternal H3K4 methylation is required for minor zygotic gene activation and early mouse embryonic development. EMBO Rep. 16, 803–812 (2015).

47. Zaret, K. S. Pioneer transcription factors initiating gene network changes. Annu. Rev. Genet. 54, 367–385 (2020).

48. Choi, S. H. et al. DUX4 recruits p300/CBP through its C-terminus and induces global H3K27 acetylation changes. Nucleic Acids Res. 44, 5161–5173 (2016).

49. Tardat, M. et al. Cbx2 targets PRC1 to constitutive heterochromatin in mouse zygotes in a parent-of-origin-dependent manner. Mol. Cell 58, 157–171 (2015).

50. Huang, B., Yang, X.-D., Zhou, M.-M., Ozato, K. & Chen, L.-F. Brd4 coactivates transcriptional activation of NF-κB via specific binding to acetylated RelA. Mol. Cell. Biol. 29, 1375–1387 (2009).

51. Zhong, H., May, M. J., Jimi, E. & Ghosh, S. The phosphorylation status of nuclear NF-κB determines its association with CBP/p300 or HDAC-1. Mol. Cell 9, 625–636 (2002).

52. Nishikimi, A., Mukai, J. & Yamada, M. Nuclear translocation of nuclear factor kappa B in early 1-cell mouse embryos. Biol. Reprod. 60, 1536–1541 (1999).

53. Hendrickson, P. G. et al. Conserved roles of mouse DUX and human DUX4 in activating cleavage-stage genes and MERVL/HERVL retrotransposons. Nat. Genet. 49, 925–934 (2017).

54. De Iaco, A. et al. DUX-family transcription factors regulate zygotic genome activation in placental mammals. Nat. Genet. 49, 941–945 (2017).

55. Vuoristo, S. et al. DUX4 regulates oocyte to embryo transition in human. bioRxiv 732289 (2020) doi:10.1101/732289.

56. Memili, E. & First, N. L. Developmental changes in RNA polymerase II in bovine oocytes, early embryos, and effect of alpha-amanitin on embryo development. Mol. Reprod. Dev. 51, 381–389 (1998).

57. Simmet, K., Zakhartchenko, V. & Wolf, E. Comparative aspects of early lineage specification events in mammalian embryos–insights from reverse genetics studies. Cell Cycle 17, 1688–1695 (2018).

58. Gerri, C., Menchero, S., Mahadevaiah, S. K., Turner, J. M. A. & Niakan, K. K. Human embryogenesis: a comparative perspective. Annu. Rev. Cell Dev. Biol. 36, 411–440 (2020).

59. Gerri, C. et al. Initiation of a conserved trophectoderm program in human, cow and mouse embryos. Nature 587, 443–447 (2020).

60. Pérez-Gómez, A., González-Brusi, L., Bermejo-Álvarez, P. & Ramos-Ibeas, P. Lineage Differentiation Markers as a Proxy for Embryo Viability in Farm Ungulates. Front. Vet. Sci. 8, 671 (2021).

61. Home, P. et al. Altered subcellular localization of transcription factor TEAD4 regulates first mammalian cell lineage commitment. Proc. Natl. Acad. Sci. 109, 7362–7367 (2012).

62. Deglincerti, A. et al. Self-organization of the in vitro attached human embryo. Nature 533, 251–254 (2016).

63. Ralston, A. et al. Gata3 regulates trophoblast development downstream of Tead4 and in parallel to Cdx2. Development 137, 395–403 (2010).

64. Chazaud, C., Yamanaka, Y., Pawson, T. & Rossant, J. Early lineage segregation between epiblast and primitive endoderm in mouse blastocysts through the Grb2-MAPK pathway. Dev. Cell 10, 615–624 (2006).

65. Roode, M. et al. Human hypoblast formation is not dependent on FGF signalling. Dev. Biol. 361, 358–363 (2012).

66. Plusa, B., Piliszek, A., Frankenberg, S., Artus, J. & Hadjantonakis, A.-K. Distinct sequential cell behaviours direct primitive endoderm formation in the mouse blastocyst. (2008).

67. Blakeley, P. et al. Defining the three cell lineages of the human blastocyst by single-cell RNA-seq. Development 142, 3151–3165 (2015).

68. Yeo, J.-C. et al. Klf2 is an essential factor that sustains ground state pluripotency. Cell Stem Cell 14, 864–872 (2014).

69. Lea, R. A. et al. KLF17 promotes human naíve pluripotency but is not required for its establishment. Development 148, dev199378 (2021).

70. Markov, G. J. et al. AP-1 is a temporally regulated dual gatekeeper of reprogramming to pluripotency. Proc. Natl. Acad. Sci. 118, (2021).

71. Ross, P. J. & Sampaio, R. V. Epigenetic remodeling in preimplantation embryos: cows are not big mice. Anim. Reprod. 15, 204–214 (2018).

72. Graf, A. et al. Fine mapping of genome activation in bovine embryos by RNA sequencing. Proc. Natl. Acad. Sci. 111, 4139 LP – 4144 (2014).

73. Bogliotti, Y. S. et al. Transcript profiling of bovine embryos implicates specific transcription factors in the maternal-to-embryo transition. Biol. Reprod. 102, 671–679 (2020).

74. Martin, M. Cutadapt removes adapter sequences from high-throughput sequencing reads. 2011 17, (2011).

75. Li, H. & Durbin, R. Fast and accurate long-read alignment with Burrows-Wheeler transform. Bioinformatics 26, 589–595 (2010).

76. Li, H. et al. The Sequence Alignment/Map format and SAMtools. Bioinformatics 25, 2078–2079 (2009).

77. Bolger, A. M., Lohse, M. & Usadel, B. Trimmomatic: a flexible trimmer for Illumina sequence data. Bioinformatics 30, 2114–2120 (2014).

78. Dobin, A. et al. STAR: ultrafast universal RNA-seq aligner. Bioinformatics 29, 15–21 (2013).

79. Love, M. I., Huber, W. & Anders, S. Moderated estimation of fold change and dispersion for RNA-seq data with DESeq2. Genome Biol. 15, 1–21 (2014).

80. Anders, S. & Huber, W. Differential expression analysis for sequence count data. Genome Biol. 11, R106 (2010).

81. Krueger, F. & Andrews, S. R. Bismark: a flexible aligner and methylation caller for Bisulfite-Seq applications. bioinformatics 27, 1571–1572 (2011).

82. Quinlan, A. R. & Hall, I. M. BEDTools: a flexible suite of utilities for comparing genomic features. Bioinformatics 26, 841–842 (2010).

83. Ramírez, F., Dündar, F., Diehl, S., Grüning, B. A. & Manke, T. deepTools: a flexible platform for exploring deep-sequencing data. Nucleic Acids Res. 42, W187–W191 (2014).

84. Kent, W. J. et al. The human genome browser at UCSC. Genome Res. 12, 996–1006 (2002).

85. Stovner, E. B. & Sætrom, P. epic2 efficiently finds diffuse domains in ChIP-seq data. Bioinformatics 35, 4392–4393 (2019).

86. Zhang, Y. et al. Model-based analysis of ChIP-Seq (MACS). Genome Biol 9, R137 (2008).

87. Heinz, S. et al. Simple combinations of lineage-determining transcription factors prime cis-regulatory elements required for macrophage and B cell identities. Mol Cell 38, 576–589 (2010).

88. Huang, D. W., Sherman, B. T. & Lempicki, R. A. Systematic and integrative analysis of large gene lists using DAVID bioinformatics resources. Nat. Protoc. 4, 44–57 (2009).

89. Huang, D. W., Sherman, B. T. & Lempicki, R. A. Bioinformatics enrichment tools: paths toward the comprehensive functional analysis of large gene lists. Nucleic Acids Res. 37, 1–13 (2009).

90. Li, Z. et al. Identification of transcription factor binding sites using ATAC-seq. Genome Biol. 20, 45 (2019).

91. Ernst, J. & Kellis, M. ChromHMM: automating chromatin-state discovery and characterization. Nat. Methods 9, 215–216 (2012).

